# Salience-driven value construction for adaptive choice under risk

**DOI:** 10.1101/506832

**Authors:** Mehran Spitmaan, Emily Chu, Alireza Soltani

## Abstract

Decisions we face in real life are inherently risky and can result in one of many possible outcomes. However, most of what we know about choice under risk is based on studies that use options with only two possible outcomes (simple gambles), so it remains unclear how the brain constructs reward values for more complex risky options faced in real life. To address this question, we combined experimental and modeling approaches to examine choice between pairs of simple gambles and pairs of three-outcome gambles in male and female human subjects. We found that subjects evaluated individual outcomes of three-outcome gambles by multiplying functions of reward magnitude and probability. To construct the overall value of each gamble, however, most subjects differentially weighted possible outcomes based on either reward magnitude or probability. These results reveal a novel dissociation between how reward information is processed when evaluating complex gambles: valuation of each outcome is based on an integrated value whereas combination of possible outcomes relies on a single piece of reward information. We show that differential weighting of possible outcomes enabled subjects to make decisions more easily and quickly. Together, these findings reveal a plausible mechanism for how salience, in terms of possible reward magnitude or probability, can influence the construction of subjective values for complex gambles. They also point to separable neural mechanisms for how reward value controls choice and attention in order to allow for more adaptive decision making.

## Introduction

Every decision we make entails some degree of risk and uncertainty and can result in one of many possible outcomes. For example, when choosing which restaurant to go to for lunch, one needs to consider many factors such as commute time, pricing, and wait time, each of which could vary depending on traffic, food availability, and the number of other customers who have also chosen that restaurant. In order to compute the overall values for such complex options or to directly compare those options, the brain has to assign a value to each possible or relevant outcome based on reward information (e.g., expected reward and probability of a given outcome) followed by combination or direct comparison of those values. Any of these processes can be very daunting when there are multiple pieces of reward information and many possible outcomes. Therefore, to enable decision making between real-world options, the brain must rely on certain mechanisms that simplify these valuation processes to reduce mental effort and to make value-based decision making more adaptive (Payne et al., 1988).

Although prospect theory (Kahneman and Tversky, 1979), the standard model of choice under risk, has been very successful in capturing many aspects of choice (Wu and Gonzalez, 1996; Birnbaum and Navarrete, 1998; Gonzalez and Wu, 1999; Abdellaoui, 2000; Bruhin et al., 2010; Glöckner and Pachur, 2012), it fails to account for choice between gambles with more than two alternative outcomes (complex gambles). As a result, various models have been proposed to tackle valuation and choice between complex gambles, including cumulative prospect theory (Tversky and Kahneman, 1992), transfer of attention exchange (Birnbaum and Navarrete, 1998; Birnbaum, 2008), decision field theory (Busemeyer and Townsend, 1993), and salience theory of choice (Bordalo et al., 2012, 2013). Interestingly, most of these models use a “rank-dependent” strategy similar to what is proposed in cumulative prospect theory. More specifically, possible outcomes are ranked based on different variables such as reward probability, and this ranking determines the influence of each outcome on the overall value. Despite the success of these models in capturing choice between complex gambles, it is still unclear how mechanisms proposed in these models can be instantiated in the brain mainly due to high-level computations required in these models.

Here, we used a combination of experimental and modeling approaches to reveal plausible neural mechanisms underlying valuation and choice between complex gambles. In two separate experiments, we examined choice between pairs of simple gambles and pairs of three-outcome (complex) gambles within the same individuals. We also developed a large family of heuristic models based on the assumption that the value of complex gambles can be constructed by differentially combining (weighting) the value of individual outcomes using a separate attentional mechanism. Critically, this attentional mechanism allows evaluation of individual outcomes and their combination to rely on different quantities (e.g., evaluation based on expected value but differential weighting based on magnitude). We then tested these models against more complex but less plausible models by fitting subjects’ choice behavior with our and those competing models. Using this approach, we addressed two key questions regarding the construction of reward value for complex gambles: 1) how are individual outcomes of a complex gamble evaluated?; and 2) how possible outcomes are compared between two complex gambles, or equivalently, how are the values of possible outcomes combined to compute the overall value of a gamble? By using various models to fit individual subjects’ choices between simple and complex gambles, we aimed to identify additional mechanisms that contribute to the valuation of and subsequent choice between complex gambles.

## Materials and Methods

### Subjects

A total of 64 human subjects (38 females) were recruited from the Dartmouth College undergraduate student population. Subjects were compensated with money and/or “t-points,” which were extra-credit points for introductory classes in the department of Psychological and Brain Sciences at Dartmouth College. The base rate for compensation was $10/hour or 1 t-point/hour. All subjects were then additionally rewarded based on their performance, up to $15/hour. This additional performance-based compensation was always monetary. None of the subjects were excluded from our final data analyses. All experimental procedures were approved by the Dartmouth College Institutional Review Board, and informed consent was obtained from all subjects before participating in the experiment.

### Overview of the experimental paradigm

Each subject performed two tasks (simple-gamble and complex-gamble tasks) in which he/she selected between a pair of gambles on every trial and were provided with feedback. In both tasks, gambles were presented as rectangular bars divided into different portions. A portion’s color indicated the reward magnitude of that outcome, and its size signaled its probability (see below). This gamble presentation was adopted from a recent study on monkeys (Strait et al., 2014) because this design allowed us to accurately present complex gambles without using any numbers, making evaluation and decision making more intuitive. For both tasks, subjects were instructed to maximize their reward points, which later translated to monetary reward or t-points, by choosing the gamble that they believed was more likely to provide more reward points. The selected gamble was resolved following each choice according to probabilities associated with possible outcomes of the chosen gamble.

Before the beginning of each task, subjects completed a training session in which they selected between two sure gambles. These training sessions were used to familiarize subjects with the associations between four different colors (purple, magenta, green, and gray) and their corresponding reward values, which depended on the task. In both training sessions, all subjects selected the gamble with higher *EV* on over 70% of the trials, indicating that they understood the color-reward associations. For the simple-gamble task, reward values were always 0, 1, 2, and 4 points. For the complex-gamble task, 0 and 1 were used, but the other two reward values varied for each subject depending on their subjective utility (see next section). No reward (0 points) was always assigned to the gray color. The color-reward assignment remained consistent for each subject throughout both the training session and its corresponding task. The color-reward assignments, however, were randomized between subjects.

### Simple-gamble task

The simple-gamble task consists of two types of trials: 1) choice between a sure option and a simple gamble with two outcomes of either a reward larger than that of the sure option or no reward with complementary probabilities; 2) choice between two simple gambles (**Fig. 1A**). Reward magnitude and probability were represented by the color and length of the corresponding portion, respectively. Subjects evaluated a total of 63 unique gamble pairs, each of which was shown four times in a random order (total of 252 trials). To make this choice task nontrivial, the gamble pairs were constructed to be relatively similar in expected value.

**Figure 1.**
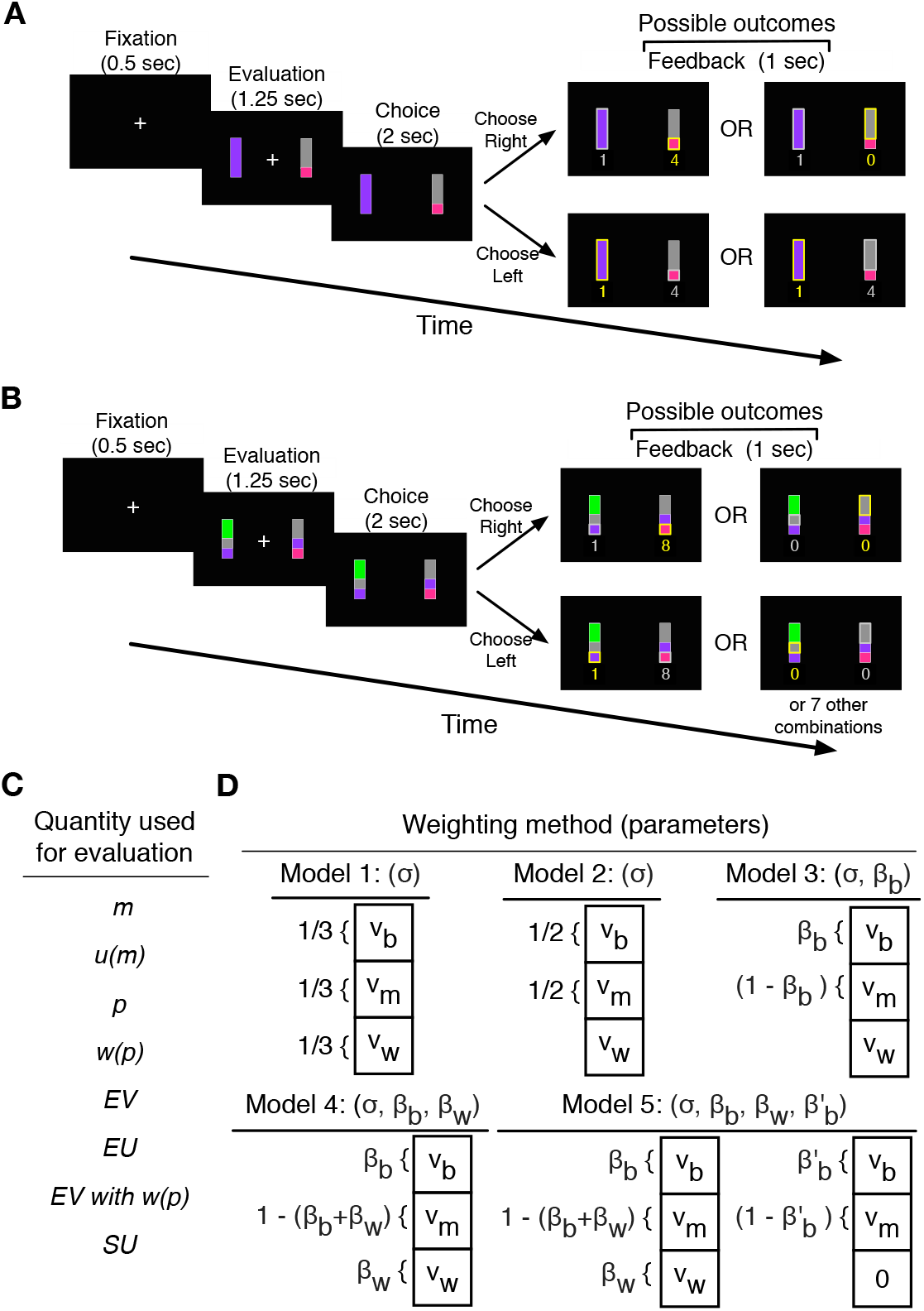
Experimental paradigm and alternative models for the construction of reward value. On each trial of the simple-gamble task (**A**) subjects selected between either a sure option and a simple gamble or a pair of simple gambles before receiving feedback. In the complex-gamble task (**B**) subjects selected between pairs of complex gambles. A simple gamble could yield reward or no reward with complementary probabilities, whereas a complex gamble offered three possible outcomes. To model the construction of reward value (**C**–**D**), we considered eight alternative quantities to *evaluate* individual gamble outcomes (**C**: *m*, magnitude; *u*(*m*), utility function; *p*, probability; *w*(*p*), probability weighting function; *EV*, expected value; *EV* with *w*(*p*); *EU*, expected utility; *SU*, subjective utility). We considered six of these quantities (*m, p, EV, EV* with *w*(*p*), *EU*, and *SU*) to *sort* possible outcomes as the best, middle, and worst (*V_b_, V_m_*, and *V_w_* correspond to the values of best, middle, and worst outcomes, respectively). The quantities *u*(*m*) and *w*(*p*) were not considered for sorting because they produce the same sorting results as *m* and *p*, respectively. We also considered five different weighting methods by which the values of individual outcomes can be combined to form the overall gamble value (**D**). The parameter *σ* represents the stochasticity in choice, which transforms the difference in reward values to the probability of choice. *β_b_* and *β_w_* are fixed model parameters denoting the differential weights associated with the best and worst outcomes, respectively. Meanwhile, *β′_b_* denotes the weight associated with the best outcome for a complex gamble that includes a zero outcome.

### Complex-gamble task

During the complex-gamble task, subjects selected between 70 unique pairs of gambles, each of which was presented four times in a random order (total of 280 trials). The complex-gamble task was similar to the simple-gamble task with a few exceptions (**Fig. 1B**). First, to make gambles that have equal subjective values during the complex-gamble task, the middle and large reward values were tailored for each subject according to their utility function estimated from their choice behavior in the simple-gamble task. More specifically, no reward (0 points) and the small reward (1 point) remained unchanged from the simple-gamble task, whereas the middle and large magnitudes were adjusted to have about double and quadruple the small reward’s utility, respectively. We kept the maximum value of reward magnitude at 10, resulting in the medians of 3 and 8 for the middle and large rewards, respectively. Although reward magnitudes associated with each color may differ between the simple-gamble and complex-gamble tasks for individual subjects, the “relative” color-reward association did not. For example, the largest reward value in the complex-gamble task would be associated with the color previously corresponding to 4 points in the simple-gamble task, etc.

Second, to increase the sensitivity of our experimental paradigm, pairs of three-outcome gambles were tailored to be very close in subjective utility for individual subjects based on their estimated utility and probability weighting functions from the simple-gamble task. Probability of each outcome was represented by the length of the corresponding portion, and to ensure that differences in probabilities were easily discernable, we constrained the gambles such that the probabilities of the three outcomes differed from one another by a value larger than or equal to 0.1. Therefore, the combination of outcome probabilities for each gamble was restricted to one of the following sets: {0.6, 0.2, 0.2}, {0.4, 0.3, 0.3}, {0.5, 0.3, 0.2}, and {0.4, 0.4, 0.2}. From the set of all possible gambles, we picked pairs of gambles for which the difference in the subjective values was less than the 5% of the difference between the maximum and minimum subjective values of all possible pairs. Our procedure and the large set of pairs guaranteed that there was no correlation between reward magnitude and probability of individual gamble outcomes (Pearson correlation: *r* = 0.01, *P* = 0.78; Spearman correlation, *r* = 0.026, *P* = 0.33). In addition, there was no significant difference between the medians of standard deviations of outcome probabilities for the best, medium, and worst outcomes. Therefore, the dispersion of outcome probabilities was not informative and could not bias subjects to attend and overweight certain outcomes. Finally, we also included 20 “catch” trials (where one of the gambles was better than the other with respect to both reward magnitude and probability) in order to detect any subject who was not attentive to the presented gambles. We found that in catch trials, all subjects selected the more valuable gamble with a probability greater than 0.7 and a median of 0.95.

### Modeling and data analysis

We fit choice behavior of individual subjects using four models to assign value to each gamble outcome. These include: expected value (EV); expected value with the probability weighting function (EV with *w*(*p*)); expected utility (EU); and subjective utility (SU). The EV model multiplied the non-zero reward magnitude (*m*) by its corresponding probability (*p*), whereas the EV with *w*(*p*) model used subjective probability *w*(*p*) instead of the actual reward probability (*m* × *w*(*p*)). The EU model assumed a nonlinear utility function, *u*(*m*), to compute the value associated with a given reward magnitude (*u*(*m*) × *p*). Finally, the SU model utilized both nonlinear utility and probability weighting functions (*u*(*m*) × *w*(*p*)).

We considered a power law for the utility function:

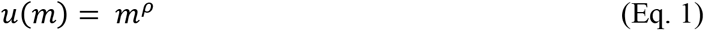

where *ρ* > 0 measures the curvature of the utility function. The probability weighting function (subjective probability) was modeled using the 1-parameter Prelec function (Prelec, 1998):

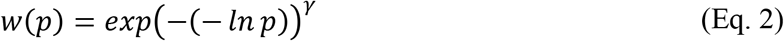

where *γ* > 0 measures the distortion of the probability weighting function. Finally, using reward values assigned to the two gambles on each trial based on a given model, the probability that the subject would choose the gamble on the right (*P_R_*) was computed as follows:

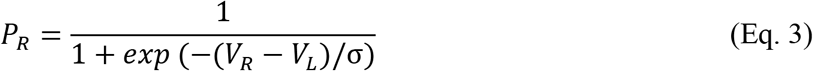

where *V_R_* and *V_L_* denote the value of the right and left gambles, respectively, and *σ* is a model parameter that measures stochasticity in choice by transforming the difference in reward values to the probability of choice.

The choice data was fit using the four aforementioned models for valuation of gambles by minimizing the negative log likelihood of the predicted choice probability given different model parameters. Minimization was done using the fminsearch function in MATLAB (Mathworks) over 50 initial model parameters. The fitting of choice behavior in the simple-gamble task allowed us to estimate the utility and probability weighting function for individual subjects. For model comparison, we used the Akaike information criterion (AIC) in order to account for the different number of parameters in different models. Although the AIC values varied between subjects, reflecting how well each subject’s choice behavior was captured, all AIC comparisons were within-subject. Therefore, variability in AIC values did not influence model comparison.

In order to further examine the quality of model comparison, we also used Vuong’s test for model selection. Specifically, Vuong test (Vuong (1989) determines the best model as the model with the log likelihood (*LL*) significantly smaller than the one of the second best model. Considering *N* samples of *LL* values for each model, Vuong statistic for comparing say models *i* and *j* is calculated using the following equation:

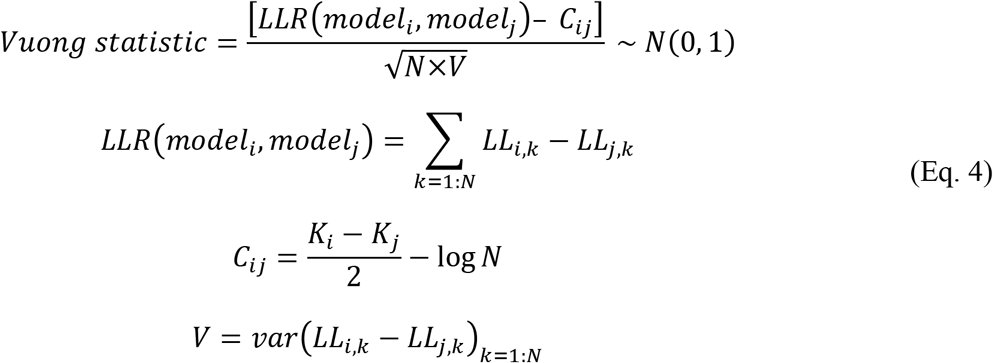

where *LLR* is the summation of *LL* ratios for the two models, *C_ij_* is a correction term for the difference of degrees of freedom between two models, *V* is the variance of *LL* ratios between two models, and *K*_1_ and *K*_2_ are the numbers of parameters in models *i* and *j*, respectively. It has been shown that Vuong statistic follows a standard normal distribution *N*(0, 1). As a result, model *i* can be considered better than *j* if Vuong statistic > 1.96 and vice versa.

We used a large set of models to fit choice behavior during the complex-gamble task. These models differed in their assumptions about how individual outcomes of a given gamble are evaluated (8 possible quantities; **Fig. 1C**), how individual outcome values are sorted (six possible quantities), and how the outcomes are combined to form the overall value of a three-outcome gamble (5 possible methods; **Fig. 1D**). Note that identical results were obtained from sorting outcomes based on *m* and *u*(*m*), and for *p* and *w*(*p*), reducing the number of sorting method to six. For the five possible weighting methods, Model 1 assumed equal weighting of all three outcomes. Models 2 and 3 ignored the worst outcome (*V_w_*) and assigned equal (Model 2) or different weights (Model 3) to the best and middle outcomes (*V_b_* and *V_m_*) when computing the overall gamble value. In contrast, Model 4 considered differential weighting spread across all three outcomes. Model 5 was similar to Model 4 when the gamble consisted of three nonzero outcomes. However, when the gamble contained a zero-reward outcome, Model 5 would not include a weight for that outcome. We also tested an additional model with equal weighting for middle and worst outcomes. The result for this model is not reported because this model was not able to successfully predict any of individual subjects. Finally, we note that although models 1, 2, and 3 can be considered as special cases of model 4, we kept these model due to smaller number of parameters in these models and possible failure of fitting procedure with larger number of parameters. All the possible combinations of assumptions resulted in the generation of 240 possible models (8 possible quantities for evaluating individual outcomes, 6 possible quantities for sorting outcomes, and 5 possible weighting methods). These models were used to fit individual subjects’ choice behavior in the complex-gamble task by utilizing a similar procedure as the one used for the simple-gamble task.

To test the ability of our fitting procedure in capturing the proposed mechanisms for the construction of overall reward value and extract the correct parameters, we simulated choice data using the proposed models over a wide range of model parameters during the same complex-gamble task performed by the subjects. We then fit the simulated data with each of these models to compute the goodness-of-fit (in terms of AIC) and to estimate the original model parameters used to simulate the data. Because there are a total of 240 possible models for this analysis, we considered a small subset of models based on the most frequent models identified in the experimental data. More specifically, we only considered three parameters for sorting the values of individual outcomes (*m, p*, and *EV*) because sorting based on the other five quantities (*u*(*m*), *w*(*p*), *EU, EV* with *w*(*p*), and *SU*) resulted in very similar outcomes as the former quantities. Moreover, we only considered Model 4 for weighting of values since this was the most extensive model used for differential weighting. Based on these specifications, we narrowed our analysis to a total of 24 models. For each of these 24 models, we generated choice data for the same complex-gamble task performed by the subjects (with the same number of parameters, trials, etc.) and fit the various choice data using each of the 24 models.

In order to quantify how easily a subject can distinguish between a pair of gambles based on their subjective values, we defined “discriminability” as follows:

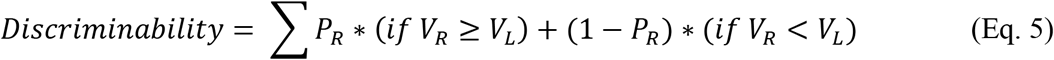

where *V_R_* and *V_L_* indicate the value of the left and right gambles, respectively. *P_R_* is the probability of selecting the right gamble, and the sum is computed over all unique pairs of gambles. The chance level of discriminability is equal to 0.5.

### Competing models of valuation and choice between complex gambles

To compare our model with existing models of valuation and choice between complex gambles, we fit choice behavior of our subjects using four different rank-dependent models. This includes cumulative prospect theory, CPT (Tversky and Kahneman, 1992), transfer of attention exchange, TAX (Birnbaum and Navarrete, 1998; Birnbaum, 2008), decision field theory, DFT (Busemeyer and Townsend, 1993), and salience theory of choice, STC (Bordalo et al., 2012, 2013). Here we provide a summary of these models and how they are implemented.

CPT generalizes prospect theory for choice under uncertainty by extending PT in multiple ways. Importantly, by adopting a cumulative representation for probability, CPT can be applied to gambles with more than two non-zero outcomes and removes the need for the editing rules of combination and dominance detection. More specifically, for gambles with strictly non-negative outcomes (*m*_1_ ≥ *m*_2_ ≥ ⋯ ≥ *m_n_* ≥ 0), the utility of gamble *G, CPT*(*G*), is equal to:

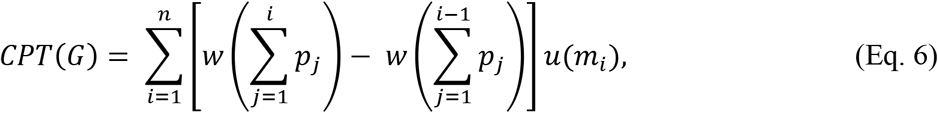

where *w*(*P_i_*) is the cumulative weighting function of (decumulative) probability of receiving more than 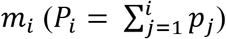, with boundary conditions of *w*(0) = 0 and *w*(1) = 1. We used the following form of the cumulative weighting function:

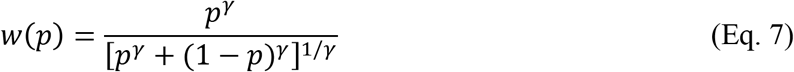

where *γ* is a free parameter. Values of *γ* > 1 and *γ* < 1 correspond to S-shaped and inverted-S-shaped curves, respectively.

In TAX, the utility of a gamble is equal to a weighted average of the utilities of the outcomes (Birnbaum and Navarrete, 1998; Birnbaum, 2008). These weights depend on the probability and rank of the branches and therefore, the relationship between branches (the so-called ‘configural’ weight). This model assumes that a decision-maker deliberates by attending to possible outcomes of an action depending on their risk attitude. Not only can branches leading to larger reward attract more attention but branches leading to lower-value outcomes can also attract greater attention if a person is risk-averse. Importantly, the weights of branches result from transfers of attention from one branch to another. If there were no configural effects, then each branch would have weights purely as a function of outcome probability, *w*(*p*). However, depending on the subject’s point of view (i.e., risk attitude), weight is transferred from branch to branch. For example, for a risk-averse subject weight can be transferred from a higher-value branch *k* to a lower-value branch *i* (*m_k_* ≥ *m_i_*). If *ω*(*p_i_, p_k_, n*) represents the weight transferred from branch *k* to branch *i*, the value of gamble *G* in TAX can then be written as:

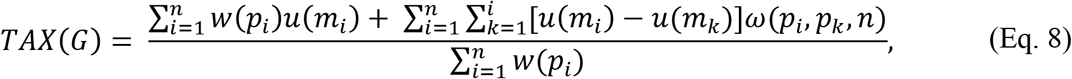

where *w*(.) the probability weighting function defined in Eq. 7 and *ω*(.) is equal to:

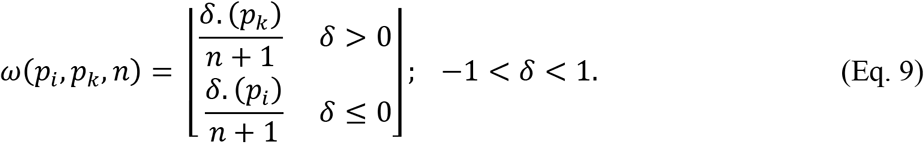

indicating that the weight transferred is a fixed proportion of the weight of the branch giving up weight. This formulation represent a general case, but assuming lower-value branches receive greater weight (*δ* > 0), a special TAX model can be written for the value of three-outcomes gambles, *G* = (*m*_1_, *p*_1_; *m*_2_, *p*_2_; *m*_3_, *p*_3_), where *m*_1_ ≥ *m*_2_ ≥ *m*_3_ ≥ 0, as follows:

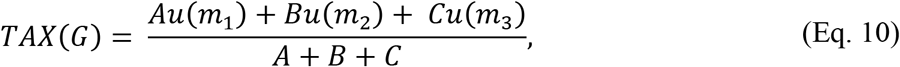

where

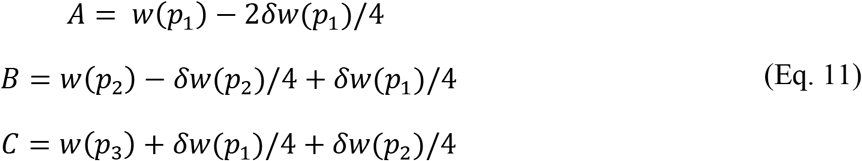

Busemeyer and Townsend (1993) derive DFT from an intuitive but sophisticated computational logic. Suppose that a decision maker attempts to choose according to rank-dependent values of alternative gambles (such as those given by CPT) but does not have an algorithm for effortlessly and quickly multiplying utilities and weights together. The decision-maker could instead proceed by sampling the possible utilities of options in proportion to their decision weights, computing the running sums of these sampled utilities for each option, and stopping (and choose) when the difference between the sums exceeds some threshold determined by the cost of sampling. Considering this algorithm, the probability of choosing an option (say the left option) based on the difference in values of the left and right options in DFT can be simplified as follows:

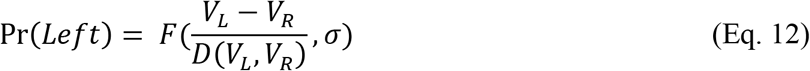

where *V_L_* and *V_R_* are the values of left and right options, respectively, and

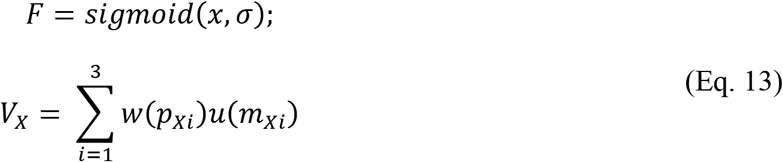

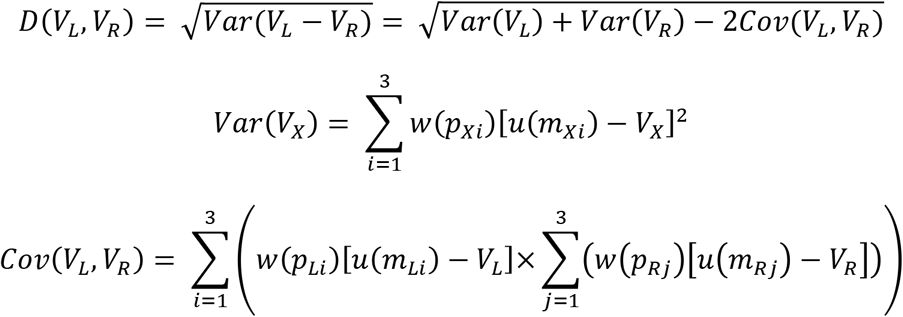

Therefore, in DFT, the difference in gamble values is normalized by a sampling process that biases choice toward gamble outcomes with extreme attribute values.

In STC, the decision-maker’s attention is drawn to (precisely defined) salient payoffs (Bordalo et al., 2012, 2013). This leads the decision-maker to a context-dependent representation of gambles in which true probabilities are replaced by decision weights distorted in favor of salient payoffs. By specifying decision weights as a function of payoffs, STC provides a unified account of many empirical phenomena, including frequent risk-seeking behavior, invariance failures such as the Allais paradox, and preference reversals. The value of a gamble in STC is computed by weighting possible outcomes based on their salience as follows (for gambles with three outcomes, *G* = (*m*_1_, *p*_1_; *m*_2_, *p*_2_; *m*_3_, *p*_3_)):

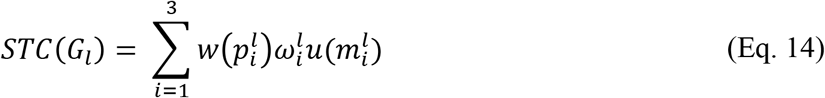

where 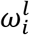 is “salient weight” for outcome *i* of gamble *l*. The salient weight for each outcome is computed using the salient ranking of each outcome:

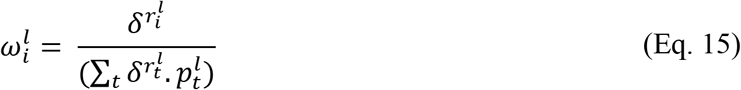

where *δ* ∈ (0,1] is a free parameter and 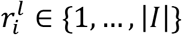 is the salient ranking of outcome i in gamble *G_l_* (lower 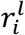 indicates higher salience). Given two outcomes of gamble *G_l_, i, ĩ* ∈ *I*, outcome *i* is considered more salient than outcome *ĩ* if 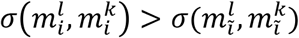 where 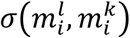 is the salient function measuring the saliency based on the outcome magnitude in the two competing gambles *l* and *k*:

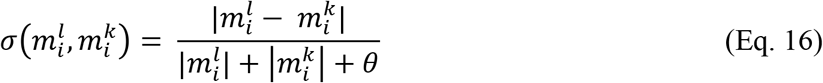

where *θ* > 0 is a free parameter.

### Our model

In our model, the probability selecting the left option is calculated based on the difference in values of the left and right options (*V_L_* and *V_R_*) as follows:

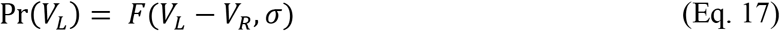

where *F* is the sigmoid function (see Eq. 13). The value of an option depends on the evaluation function, sorting algorithm, and the weighting model:

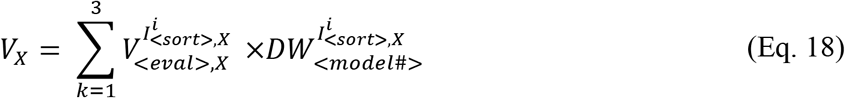

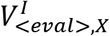 is the value of outcome *i* of option *X* based on a given evaluation function, *eval_func_*,

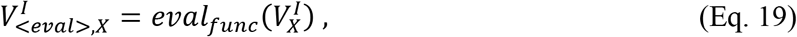

which can be equal to magnitude (*m*), utility function (*u*(*m*)), probability (*p*), weighted probability (*w*(*p*)), expected value (*m*×*p*), expected utility (*u*(*m*)×*p*), expected value with weighted probability (*m*×*w*(*p*)), or subjective utility (*u*(*m*) × *w*(*p*)). Sorting index for each outcome, 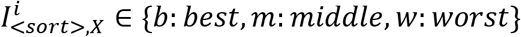 is computed after sorting outcomes based on the chosen sorting label:

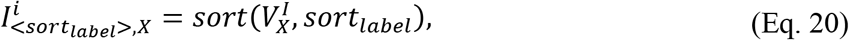

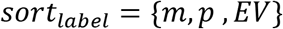

Finally, 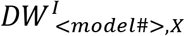 determines the value of differential weighting for each outcome based on the sorting index and one of five possible weighting models (**Fig. 1D**). Model 1 equally weights the three possible outcomes, 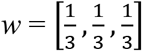, whereas Model 2 assigns equal weights to the two best outcomes and ignore the worst outcome, 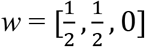. Model 3 uses a single parameter to distribute weights between two best outcomes while assigning zero weight to the worst outcome, *w* =[*β_b_*, (1 − *β_b_*), 0]. Model 4 distributes weights to the three outcomes using two parameters, *w* = [*β_b_*, 1 − (*β_b_* + *β_w_*), *β_w_*]. Finally, Model 5 is similar to Model 4 but uses a separate parameter for distributing weight when there is a zero-magnitude outcome.

## Results

The experiment consisted of two sessions. In the first session, subjects selected between either a sure gamble and a simple gamble or between a pair of simple gambles (simple-gamble task; **Fig. 1A**). In the second session, however, decisions were made between pairs of three-outcome gambles (complex-gamble task; **Fig. 1B**). In both tasks, gambles were presented as rectangular bars divided into different portions. A portion’s color indicated the reward magnitude of that outcome, and its size signaled its probability (see Materials and Methods).

To examine whether subjects comprehended the objective of both tasks, we computed the probability of selecting the gamble with a larger expected value (*EV*) on a given trial for each subject. During the simple-gamble task, subjects selected the gamble with a higher *EV* more often with a median equal to 0.79 across all subjects. This tendency was weaker in the complex-gamble task with the median equal to 0.57. Less frequent selection of gambles with a higher *EV* (i.e. more noisy behavior) was expected during the complex-gamble task since the pairs of presented gambles were closer in *EV* in this task. However, on “catch” trials of the complex-gamble task (where one of the gambles was better than the other with respect to both reward magnitude and probability), subjects selected the better option with a probability larger than 0.7 and a median of 0.95.

We then used various models to fit choice behavior in order to identify how individual subjects constructed the overall value of gambles in each task. For the simple-gamble task, we considered four models for evaluation of gambles: expected value (EV) equal to the product of reward probability and magnitude; expected value with the probability weighting function (EV with *w*(*p*)) based on the objective value for magnitude and a weighting function for reward probability; expected utility (EU) based on the objective value for probability and a utility function (*u*(*m*)) for magnitude; and subjective utility (SU) based on utility and probability weighting functions (see Materials and Methods). In contrast to simple gambles with only one non-zero reward outcome, complex gambles had up to three non-zero outcomes. Therefore, we used a large family of models to examine how subjects evaluated the latter gambles. These models differed in their assumptions about the quantity used for the *evaluation* of individual outcomes of a given gamble (eight possible quantities; **Fig. 1C**), the quantity used for *sorting* and in turn assigning different weights to individual outcomes (six possible quantities), and how their outcome values are combined to construct the overall value of a complex gamble (5 possible methods; **Fig. 1D**), resulting in a total of 240 unique models (see Materials and Methods). For model comparison, we used the Akaike information criterion (AIC) to account for different numbers of parameters in different models. Similar results were obtained when using the Bayesian information criterion (BIC).

Considering the complexity of the proposed models, we first tested whether our fitting procedure can identify the underlying mechanisms for the construction of overall reward value and estimate the associated parameters correctly. More specifically, we simulated choice data using the proposed models over a wide range of model parameters for the same complex-gamble task performed by the subjects. We subsequently fit the simulated data with each of these models to compute the goodness-of-fit (in terms of AIC) and to estimate the original model parameters used to simulate the data (see Materials and Methods for more details). We found that for most cases, the model used to generate the data provided the best fit, indicating that the fitting could be used to identify the mechanism underlying valuation (**Fig. 2**). However, the fitting procedure could not perfectly distinguish between certain quantities used for the evaluation of individual outcomes: *m* and *u*(*m*); *p* and *w*(*p*); and *EV, EV* with *w*(*p*), *EU*, and *SU*. As a result, we took caution when interpreting the fitting results of actual subjects’ data with regards to the quantities used to evaluate individual outcomes. In addition, we found that the average AIC for fitting data generated with single-attribute evaluation methods was only slightly worse for the combined-attribute than the single-attribute evaluation methods, when sorting was preformed based on magnitude (**Fig. 2-1A-C**). However, this resulted in misidentifying the single-attribute evaluation methods as combined-attribute ones in only 15% of the instances across a large set of model parameters (**Fig. 2-1D-E**). Finally, the overall relative differences between the actual parameters and the parameters estimated by the same model used for generating the data were very small and showed little to no systematic biases (**Fig. 2-2**). Overall, these results support the feasibility of our fitting approach for identifying mechanisms used for the construction of overall reward value in the real data.

**Figure 2.**
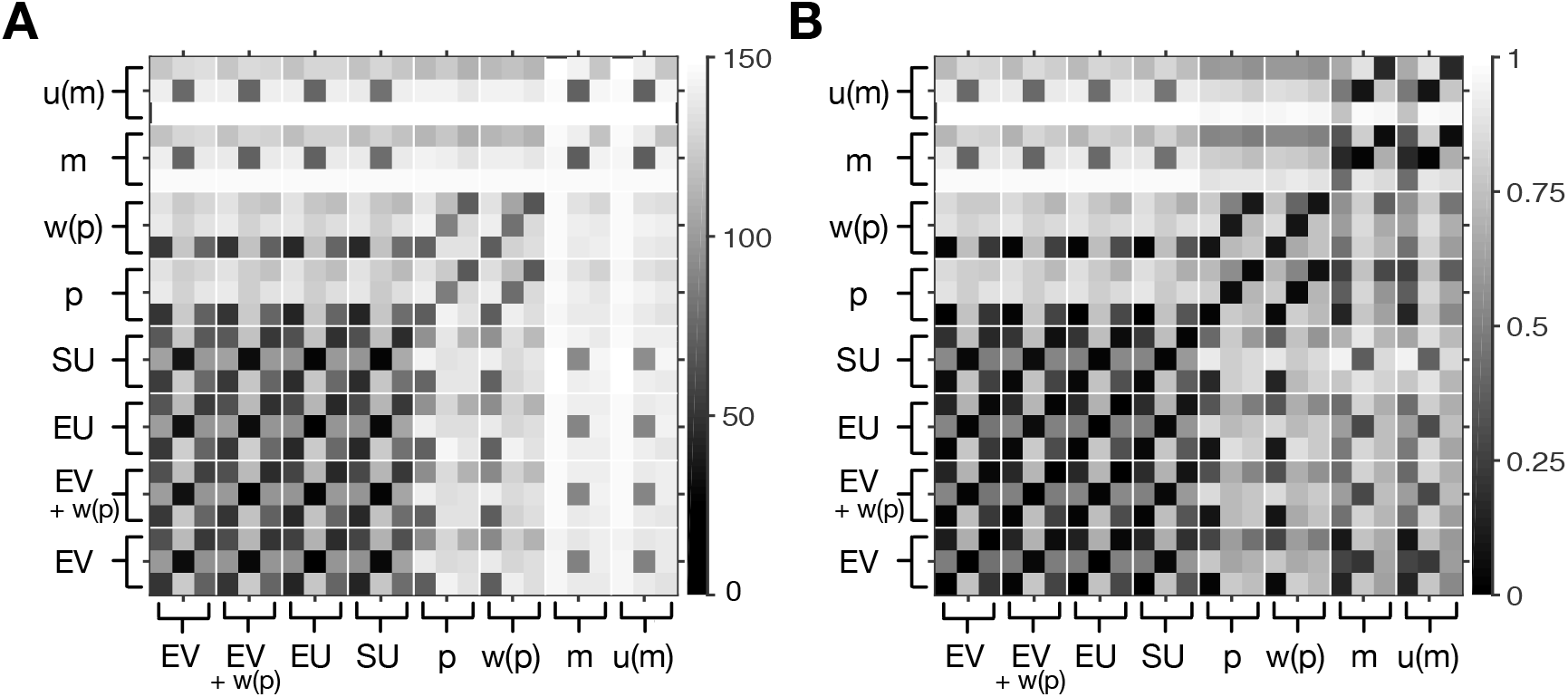
Fitting method was able to correctly identify the quantity used for evaluating individual gamble outcomes and for sorting. (**A**) Plot shows the goodness-of-fit (in terms of the average AIC over a set parameters) for fitting choice data generated with a given model and fit with the same or different models (total 24 models). The models used to generate and fit the data are indicated on the x- and y-axis, respectively. For each quantity used for evaluating individual outcomes, there are three ways to sort based on magnitude, probability, and *EV* (first, second, and third elements; see Materials and Methods). In most cases, the model used to generate the data provided the best fit for the generated data, as reflected by the darkest square in each column being on the diagonal. However, the fitting procedure could not perfectly distinguish between certain quantities used for generating the value of individual gamble outcomes: *m* and *u*(*m*); *p* and *w*(*p*); and *EV, EV* with *w*(*p*), *EU*, and *SU*. (**B**) The same as in panel A but when the AIC values were normalized by the maximum value in each column.

**Figure 2-1.**
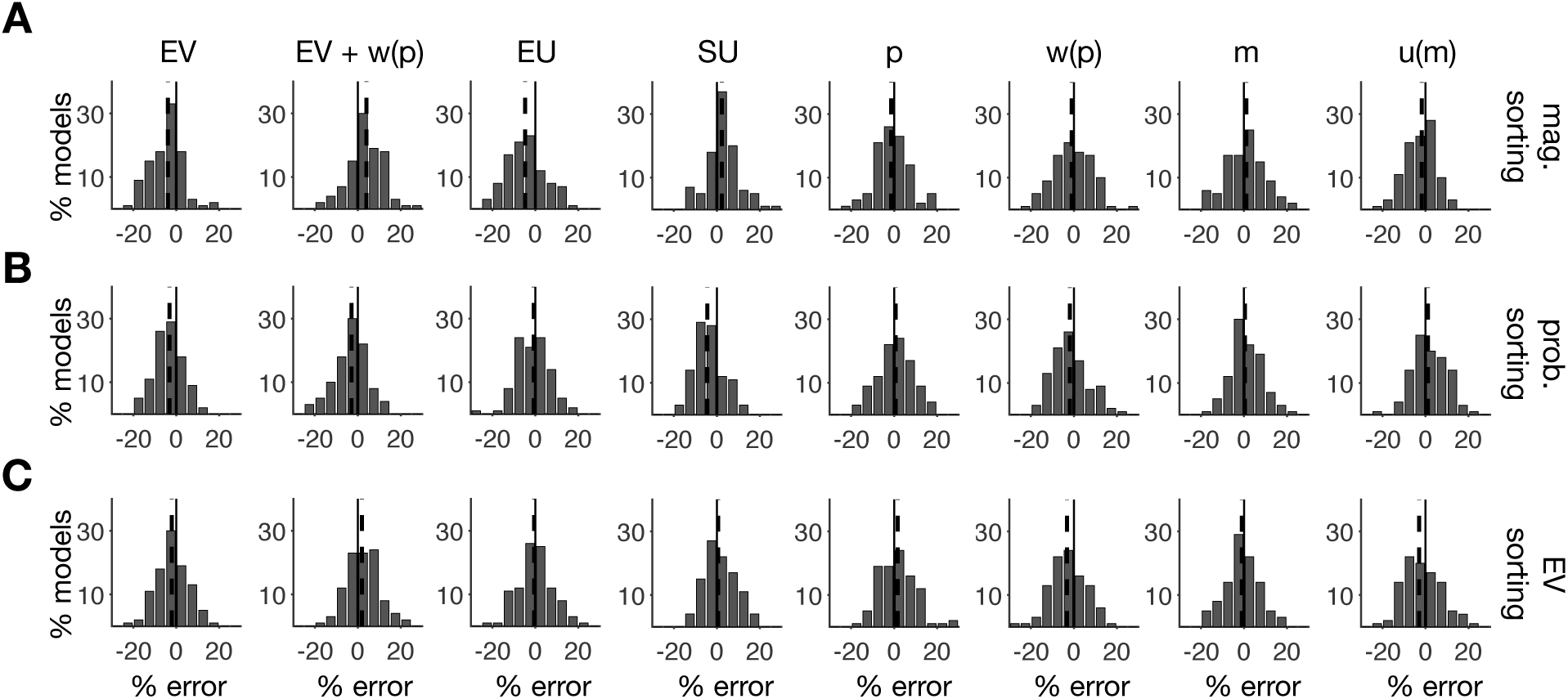
Small error in estimating model parameters based on the model used to generate the choice data. Each column corresponds to a different quantity used to evaluate individual gamble outcomes, and each row corresponds to a different quantity used for sorting outcomes (magnitude, probability, and *EV*), which were used to assign different weights to each outcome. Overall, there were small error and negligible bias in the estimated parameters.

**Figure 2-2.**
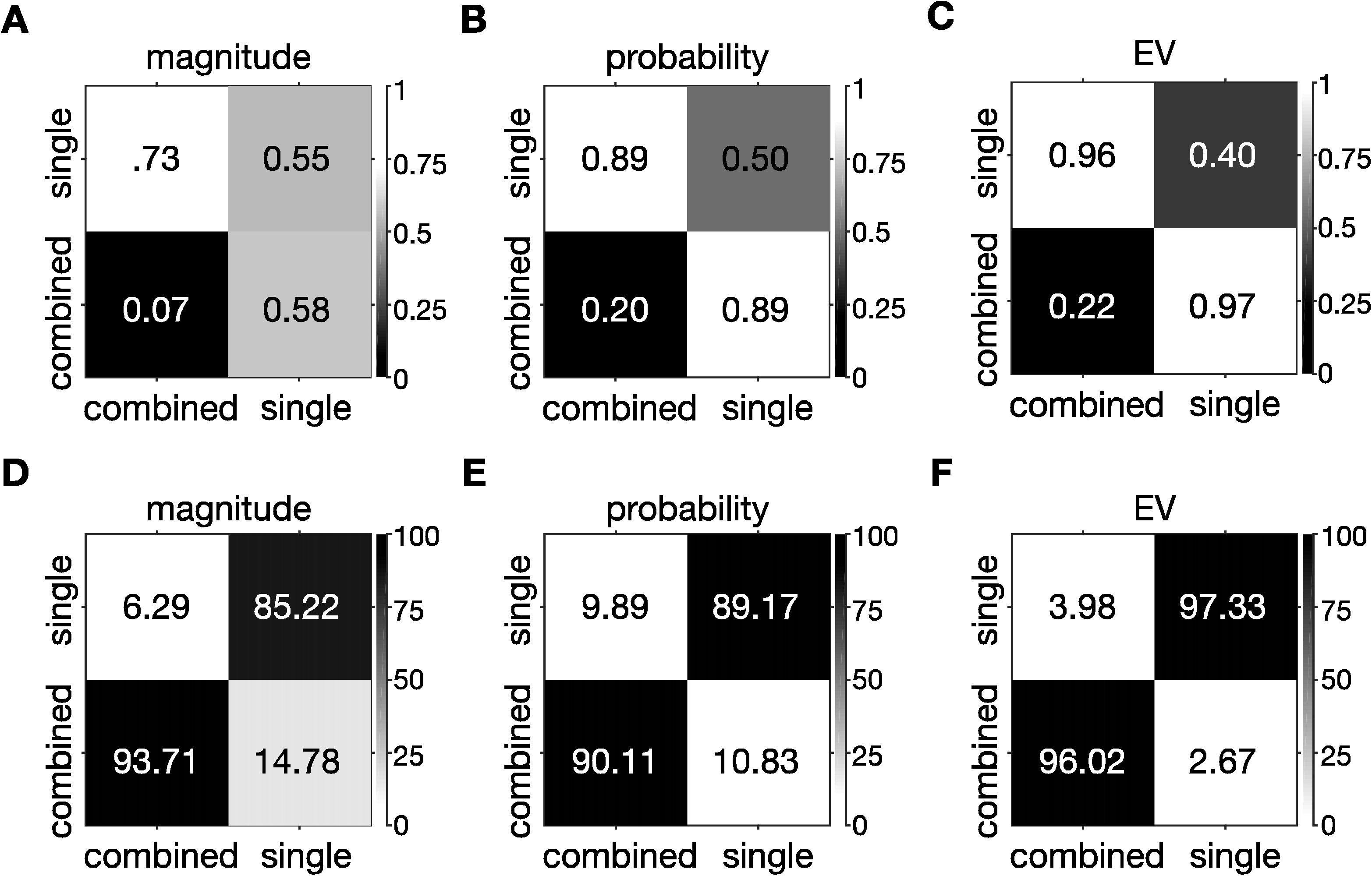
Fitting method was able to correctly identify the quantity used for evaluating individual gamble outcomes and for sorting. (**A-C**) Plot shows the goodness-of-fit (in terms of the average normalized AIC over a set parameters) for fitting choice data generated with a given model and fit with the same or different models. The models used to generate and fit the data are indicated on the x- and y-axis, respectively. Each column shows average over a type of sorting method used for evaluating individual outcomes (single: sorting based on *m, u*(*m*), *p, w*(*p*); combined: sorting based on *EV, EV*+*w*(*p*), *EU, SU*). (**D-F**) The same as in panel A-C but showing the percentage of models fit into each category.

### Evaluation of simple gambles and individual outcomes of complex gambles conformed to PT

Fitting of choice behavior showed that the SU model provided the best fit for the majority (83%) of subjects in the simple-gamble task (**Fig. 3A**). These results indicate that subjects mainly used subjective utility to evaluate simple gambles. The estimated utility and probability weighting functions conformed to the predictions of PT. More specifically, the majority of subjects (56 out of 64, corresponding to ~88% of subjects) exhibited concave utility functions (**Fig. 3B**), and most subjects (43 out of 64, corresponding to ~67% of subjects) showed inverse-S-shaped probability distortion (**Fig. 3C**). Overall, these results demonstrate that PT can successfully account for choice between simple gambles.

**Figure 3.**
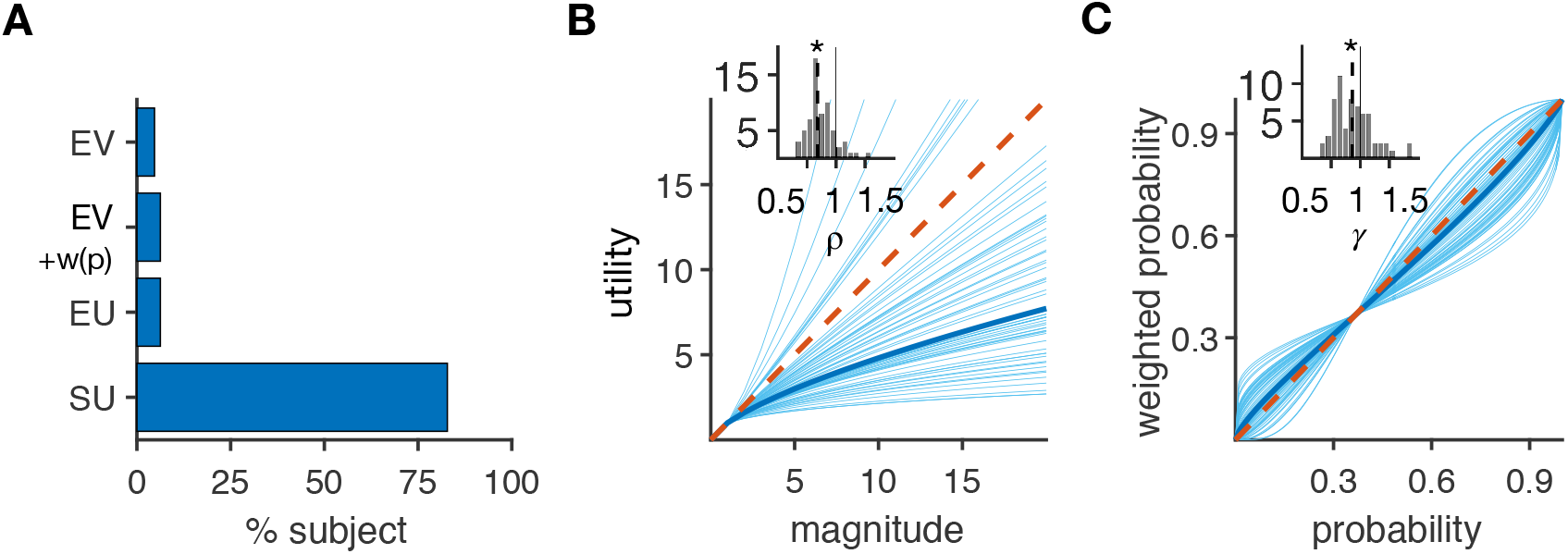
Prospect theory captures choice between simple gambles in most subjects. (**A**) Plot shows the fraction of subjects for whom a given model provided the best fit. The SU model provided the best fit for the majority (53 out of 64) of subjects. (**B**) Plotted is the estimated utility function for individual subjects (light blue curves) based on the SU model. The dark blue curve shows a utility function formed based on the median of the distribution of estimated exponents of the utility function (*ρ*), as shown in the inset. The red dashed line shows the unity line. The solid and dashed lines in the inset indicate *ρ* = 1 and the median of *ρ* values (0.38<*ρ*<1.53 median: *ρ*=0.63), respectively; the asterisk shows the median is significantly different from 1 (two-sided sign-test, *P* = 2.01 ×10^−34^, *d* = 3.10). (**C**) Plotted is the estimated probability weighting function for individual subjects based on the SU model. The dark blue curve shows a probability weighting function formed based on the median of the distribution of estimated exponents of the probability weighting function (*γ*), as shown in the inset. The solid and dashed lines in the inset indicate *γ* = 1 and the median of *γ* (0.36< *γ* <1.89 median: *ρ*=0.88), respectively. The asterisk shows the median is significantly different from 1 (two-sided sign-test, *P* = 0.008, *d* = 2.70).

### Subjects used different quantities to evaluate and weight the outcomes of complex gambles

We used various combinations of models for sorting, evaluating, and weighting gamble outcomes in order to examine the construction of reward value for complex gambles (**Fig. 1C-D**). We first examined the quantity used by subjects to sort the possible outcomes of complex gambles into best, middle, and worst outcomes in order to differentially weigh them. We found that 61% of subjects sorted gamble outcomes based on outcomes’ reward magnitudes or probabilities (**Fig. 4A**). This percentage was larger than the percentage of subjects who used an integrated value (*EV, EV* with *w*(*p*), *EU*, or *SU*) for sorting outcomes (*χ*^2^ (1) = 5.28, *P* = 0.012).

**Figure 4.**
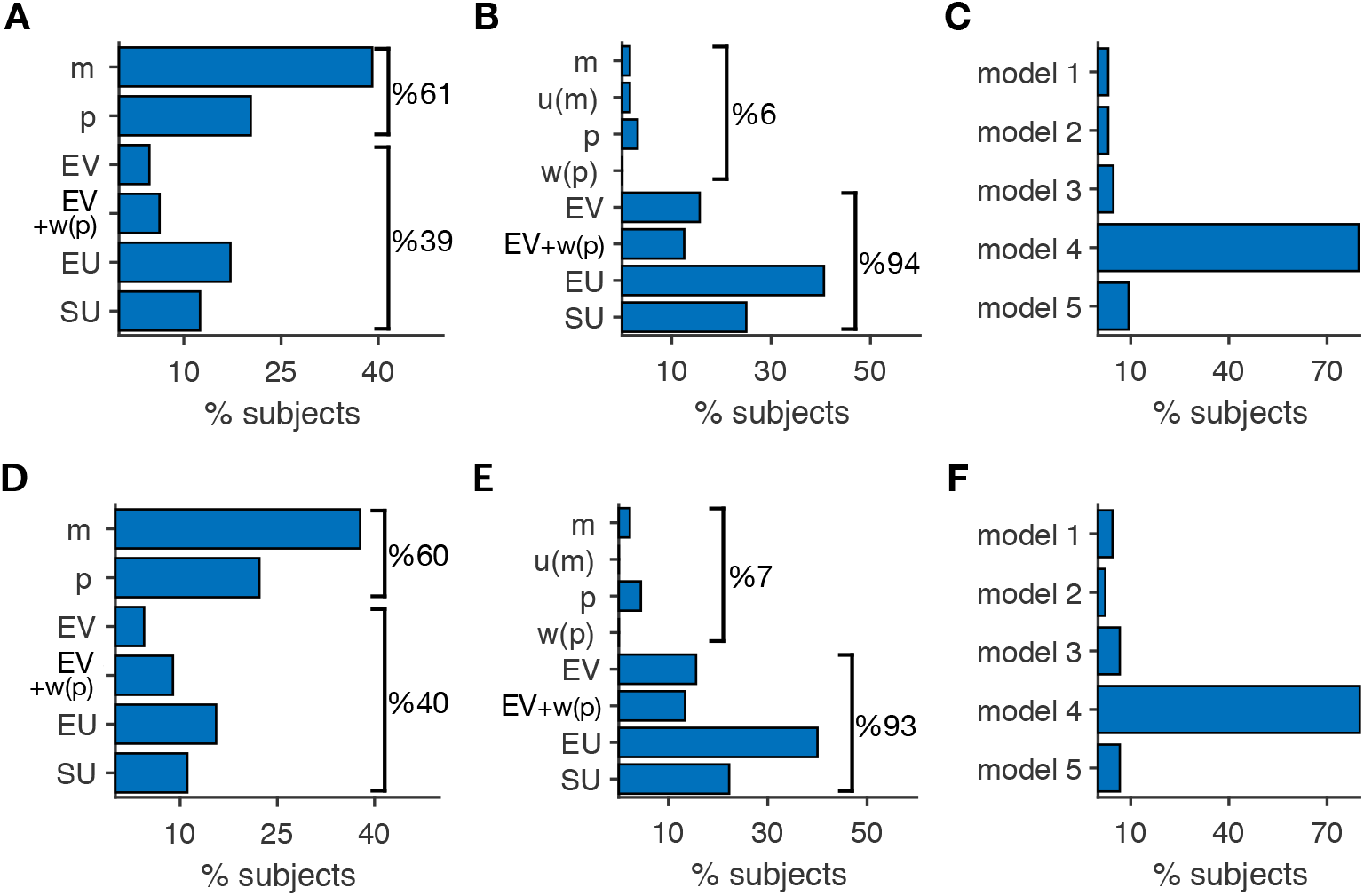
Most subjects sorted and weighted possible outcomes based on reward magnitude or probability but evaluated each gamble outcome based on an integrated value. (**A**) Plot shows the fraction of subjects whose choice was best fit by a given model for sorting. Reward magnitude was the quantity most used amongst subjects (40%) to sort gamble outcomes, whereas 21% used reward probability for sorting. Note that identical results were obtained from sorting outcomes based on *m* and *u*(*m*), and thus, results from these alternative sorting methods are combined. For the same reason, results for *p* and *w*(*p*) were also combined. (**B**) Plot shows the fraction of subjects whose choice was best fit by a given model to evaluate individual gamble outcomes. *EU* was the quantity most used by subjects (41%) to evaluate individual outcomes in complex gambles. (**C**) Plot shows the fraction of subjects whose choice was best fit by a given model of outcome weighting. Model 4 provided the best fit for the majority (51 out of 64, corresponding to 80%) of subjects. (**D–F**) Plots are similar to panels **A–C** but show the results for subset of subjects for whom the fit based on their best model was significantly different from other models (*N* = 45). 38% of subjects sorted gamble outcomes based on reward magnitude, whereas 22% used reward probability for sorting (D). 40% of subjects used *EU* to evaluate individual outcomes in complex gambles (E). Model 4 provided the best fit for the majority (36 out of 45, corresponding to 80%) of subjects (F).

We then compared these models in terms of their assumption regarding what quantity is used to evaluate individual gamble outcomes. To identify the best overall model, we computed the percentage of subjects who used a given quantity to evaluate individual outcomes, as determined by the model that provided the best fit in terms of the AIC. We found that 41%, 25%, 16%, and 12% of subjects evaluated individual outcomes based on *EU, SU, EV*, and *EV* with *w*(*p*), respectively (**Fig. 4B**). Therefore, these results illustrate that most subjects (60 out of 64, corresponding to 94% of subjects) integrated information about reward probability and magnitude to evaluate individual outcomes of complex gambles.

As mentioned earlier, our method could misidentify single-attribute evaluation methods with combined-attribute ones in ~15% instances, when sorting based on magnitude. However, a small number of subjects (4 out of 64) whose choice behavior was better fit with an evaluation based on a single piece of information indicates that the aforementioned misidentification was probably small and the vast majority of our subjects evaluated individual gamble outcome using a combination of information. In addition, the fitting procedure could misidentify the exact evaluation method among combined-attribute methods (i.e., *EV, EV* with *w*(*p*), *EU*, and *SU* methods). Therefore, caution should be used in interpreting the exact method used for combined-attribute evaluation.

Our results also show that a smaller fraction of subjects used a nonlinear probability function for the evaluation of individual outcomes of complex gambles than for simple gambles (complex gambles, 37%; simple gambles, 89%; *χ*^2^ (1) = 34.43, *P* = 4.4×10^−10^ *N* = 64). Therefore, although 94% of subjects used an integrated quantity to evaluate individual outcomes, most subjects used a single piece of reward information when sorting outcomes for weighting. This stark difference demonstrates that separate mechanisms were involved in evaluating and combining the values of individual outcomes in order to construct an overall value for complex gambles.

To determine how subjects combined outcome values to form the overall value of a gamble (or equivalently, to directly compare gambles), we then compared five alternative models of outcome weighting (**Fig. 1D**). This analysis revealed that Model 4, which assigned different weights to the three possible outcomes, provided the best fit for a majority (80%) of the subjects (**Fig. 4C**). Moreover, choice behavior of only 5 out of 64 subjects (8%) were best fit by Models 2 and 3, indicating that the majority of subjects (92%) considered the values of all three possible gamble outcomes when making decisions. We found similar dissociation between the quantities used for sorting possible outcomes and for combining them based on the mean AIC across all subjects instead of the best model for individual subjects (**Fig. 4-1**).

**Figure 4-1.**
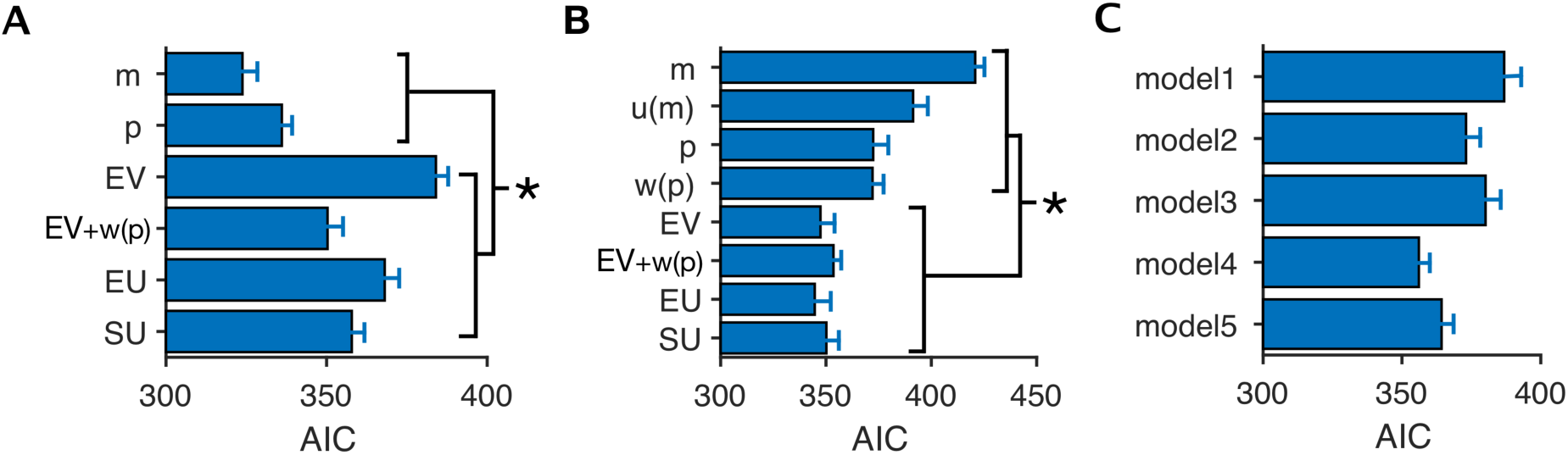
Models with integrated outcome values can better predict choice behavior. Plots show mean and standard error of AIC values over all subjects for all the models with different quantities for sorting (**A**), valuation of individual outcomes (**B**), and different types of models (**C**). We calculated the mean of all the AICs coming from the models in each category for all subjects to calculate a single mean value for that category. To determine the significance, we calculated the difference between AIC values of all the models with integrated outcome values versus all the models with individual outcome values for sorting (**A**) or valuation (**B**). The asterisk indicates that the median difference was statistically different than zero (two-sided sign-test, *P* < 0.05).

To examine whether the best model for each subject is significantly better than the rest of the models, we performed Vuong test (Vuong, 1989) (see Materials and Methods). We found that for 19 subjects (~30% of subject) there was no significant difference between the best model (model with minimum log likelihood value) and the second best model. For the remaining majority of subjects (45 equal to ~70% of subjects), however, we found the same pattern of results as in our original analysis using all data (**Fig. 4D-F**).

In our study, reward magnitude was represented by specific combinations of colors for different subjects. Therefore, we also examined that our observations were not due to specific combinations of color-reward assignments (e.g., red for the largest reward could be more effective than green); that is, there was no systematic color bias. We categorized subjects into six possible groups based on their color-reward assignments and examined differential weighting for each group (**Fig. 4-2**). This analysis revealed no systematic differences between the groups, indicating that differential weighting of possible outcomes was not due to specific color-reward assignments.

**Figure 4-2.**
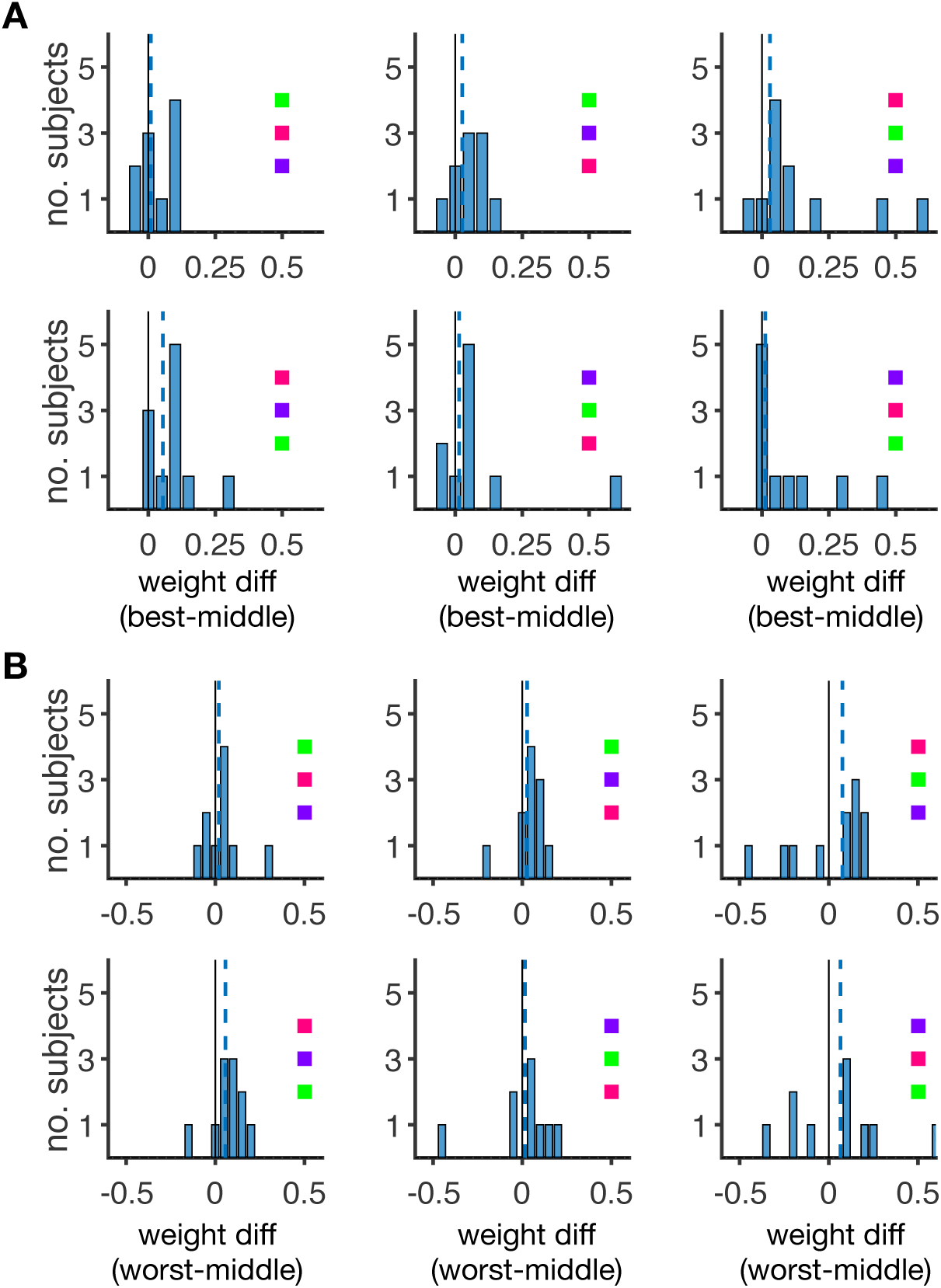
Differential weighting was not due to specific color-reward assignments. Plots show the distribution of relative weight differences separately for each group of subjects with a specific color-reward assignment. Each inset indicates the color-reward assignment specific to the corresponding group, and colors associated with the largest, middle, and smallest reward magnitudes are shown from top to bottom. There were no significant weight differences between the best and middle outcomes (**A**) or the worst and middle outcomes (**B**) (two-sided sign-test; *P* > 0.05). The blue dashed lines indicate the medians.

### Differential weighting could not be captured by PT and captured choice behavior better than competing models

Considering the complexity of models used for fitting, we also examined whether the proposed differential weighting is necessary to capture choice behavior (i.e. our results are not affected by over-fitting), and whether our fitting approach could identify the underlying model parameters without any systematic bias in the presence and absence of differential weighting (the latter was a more specific case of the analysis presented in **Fig. 2**). Therefore, we generated choice data based on three models that evaluated individual gamble outcomes using subjective value (the most common quantity used for evaluation amongst our subjects), sorted these outcome values based on magnitude, probability, or expected value, and then combined these outcome values based on Model 4 (the most common weighting method used amongst our subjects) using a wide range of model parameters. We then fit these simulated data with each of the models used to generate the data as well as with the model without differential weighting (Model 1).

We found that the model with differential weighting used to generate the data was able to fit simulated choice data very well (**Fig. 4-3** and **Fig. 4-5**) and captured the original model parameters without any bias and with small error in most cases (**Fig. 4-4** and **Fig. 4-6**). In contrast, the model without differential weighting was not able to fit the simulated choice data well and provided systematically-biased estimates of risk-preference parameters. More specifically, the estimated *ρ* and *γ* values based on the model without differential weighting were smaller than the actual values, indicating that some of the observed concavity of the utility function and curvature of the inverse-S-shaped probability weighting function could be due to differential weighting. Together, these results not only validate our approach but also illustrate that prospect theory cannot be used to fit data for which value construction is influenced by differential weighting.

**Figure 4-3.**
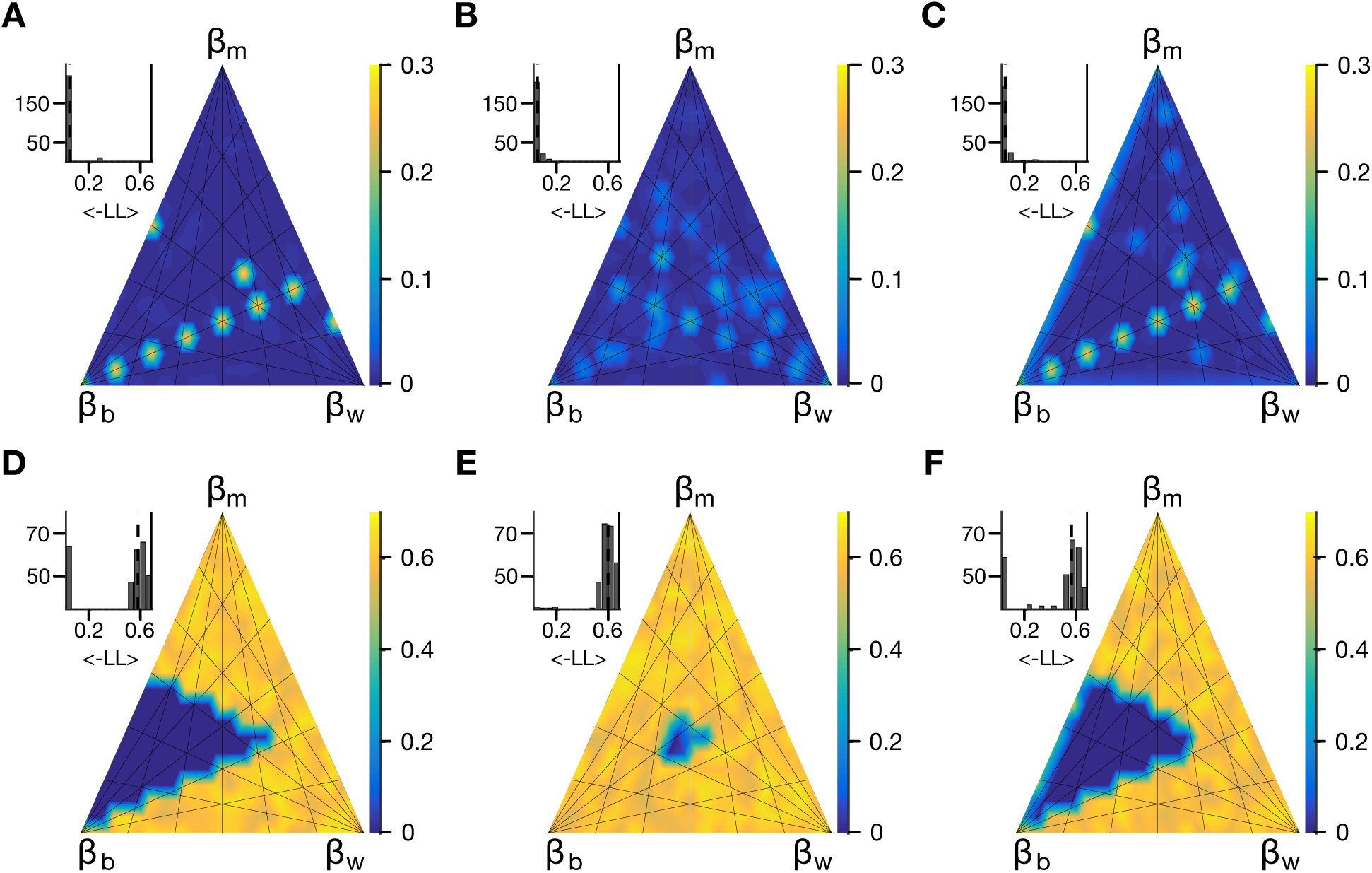
The model without differential weighting cannot fit choice data generated with differential weighting and linear utility and probability weighting functions. (**A**) Plot shows the average goodness-of-fit (in terms of negative log likelihood; a smaller number corresponds to a better fit) for fitting choice data generated by models with different sets of weights, as indicated by distance from each corner of the ternary plot, and with sorting based on reward magnitude. For these simulations, the values of individual gamble outcomes were computed using linear utility and probability weighting functions (*ρ* = 1.0 and *γ* = 1.0). The inset shows the distribution of average negative log likelihood across all models. (**B-C**) The same as in panel A but for sorting based on reward probability and *EV*, respectively. (**D-F**) The same as in panels A-C but when the choice data are fit with the model without differential weighting. Overall, the model without differential weighting failed to fit the data for most weight values.

**Figure 4-4.**
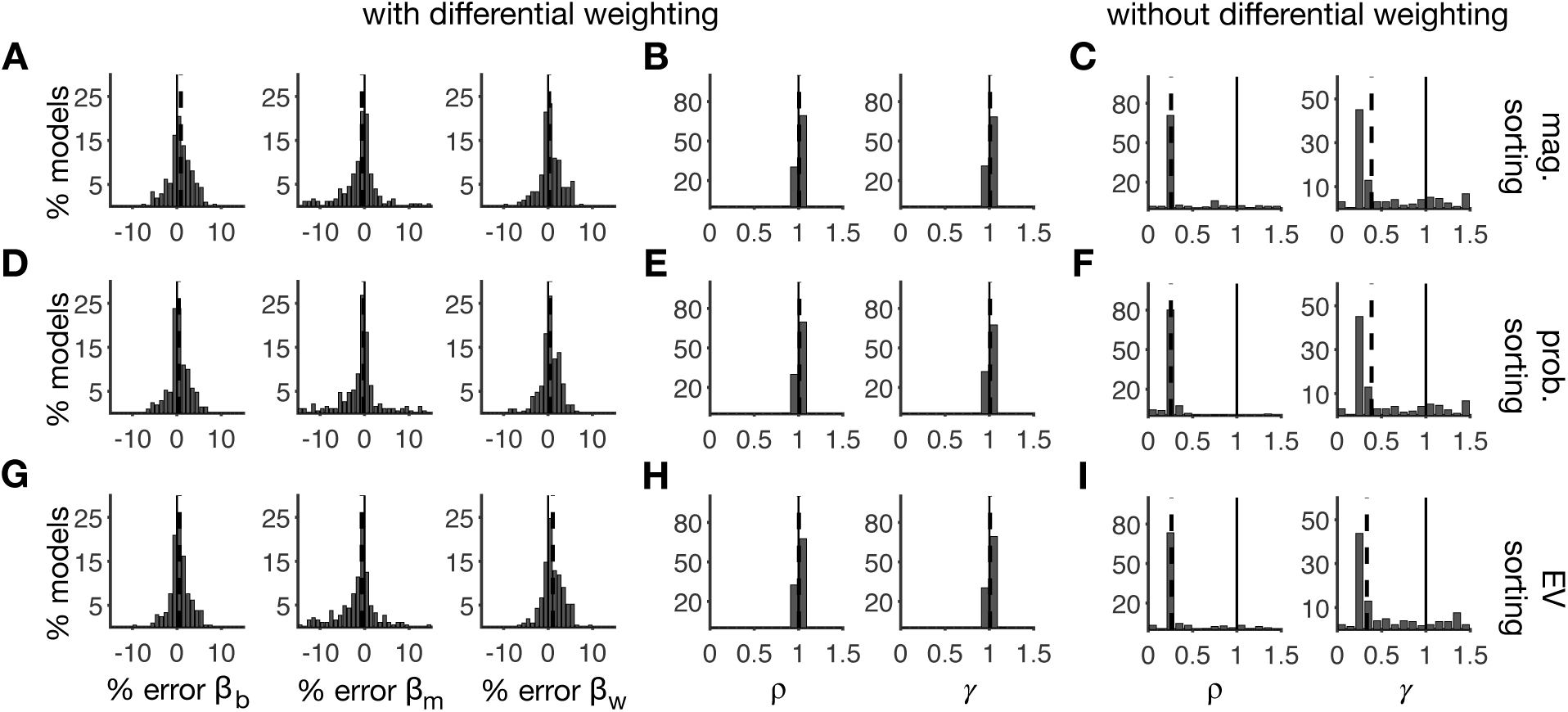
The model with differential weighting can estimate original parameters without a significant bias, whereas the model without differential weighting systematically underestimates *ρ* and *γ* for the data generated with differential weighting and linear utility and probability weighting functions. (**A**) Plots show the distribution of the relative error in the estimation of weights assigned to the best, medium, and worst outcomes (from left to right). For these simulations, the values of individual gambles were computed using linear utility and probability weighting functions (*ρ* = 1.0 and *γ* = 1.0), and outcomes were sorted based on reward magnitude. (**B**) The distribution of estimated values for *ρ* and *γ* across all models. Overall, the estimated parameters were centered around the actual values. (**C**) The distribution of estimated values for *ρ* and *γ* when choice data are fit with a model without differential weighting. The estimated parameters were significantly smaller than the actual values. (**D-F**) The same as in panels A-C but for sorting based on reward probability. (**G-I**) The same in panels A-C but for sorting based on *EV*.

**Figure 4-5.**
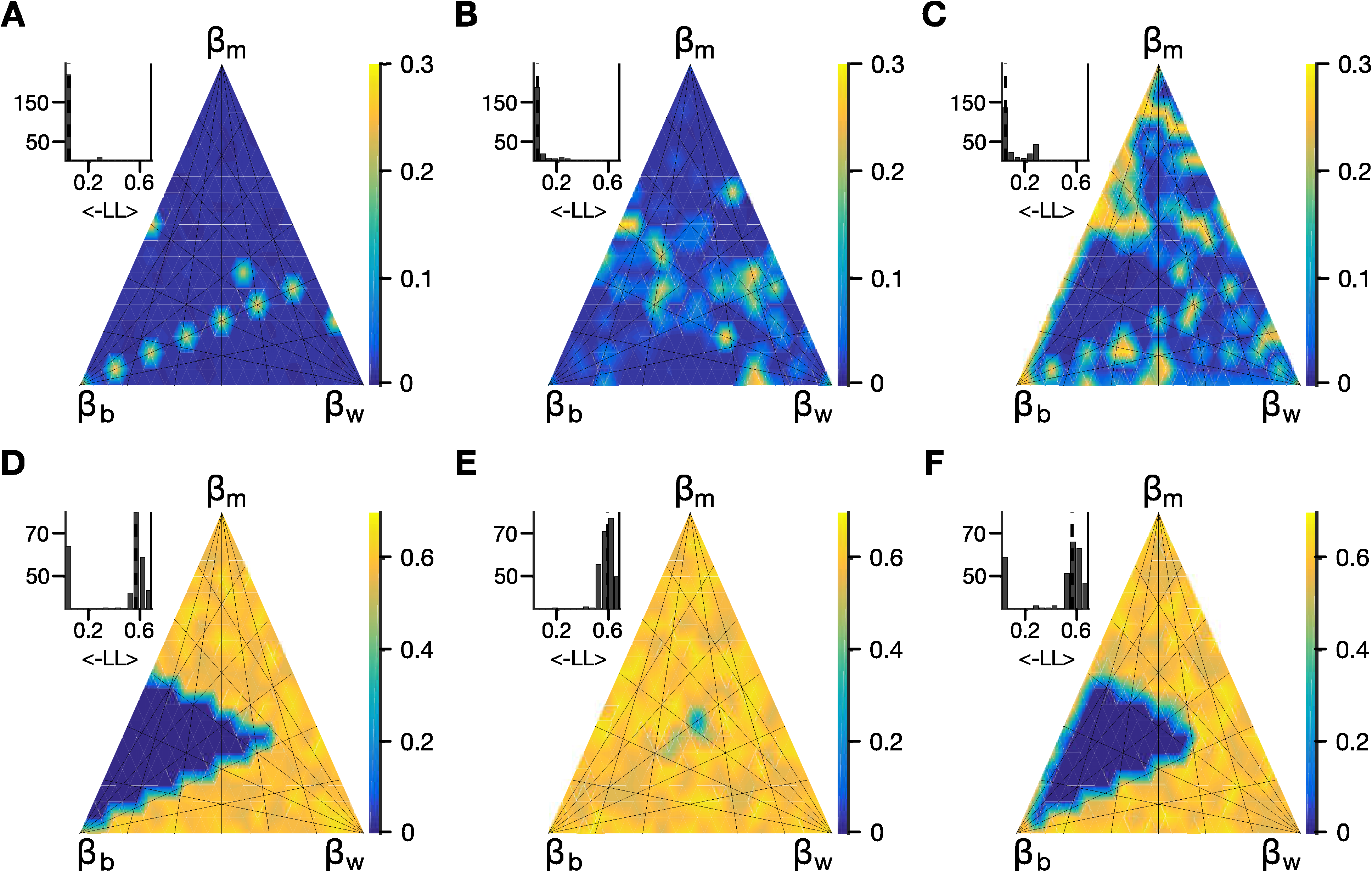
The model without differential weighting cannot fit choice data generated with differential weighting and nonlinear utility and probability weighting functions. (**A**) Plot shows the average goodness-of-ft (in terms of negative log likelihood; a smaller number corresponds to a better fit) for fitting choice data generated by models with different sets of weights, as indicated by distance from each corner of the ternary plot, and with sorting based on reward magnitude. For these simulations, the values of individual gamble outcomes were computed using a concave utility function and an inverse-S-shaped probability weighting function (*ρ* = 0.6 and *γ* = 0.9). The inset shows the distribution of average negative log likelihood across all models. (**B-C**) The same as in panel A but for sorting based on reward probability and *EV*, respectively. (**D-F**) The same as in panels A-C but when the choice data are fit with the model without differential weighting. Overall, the model without differential weighting failed to fit data for most weight values.

**Figure 4-6.**
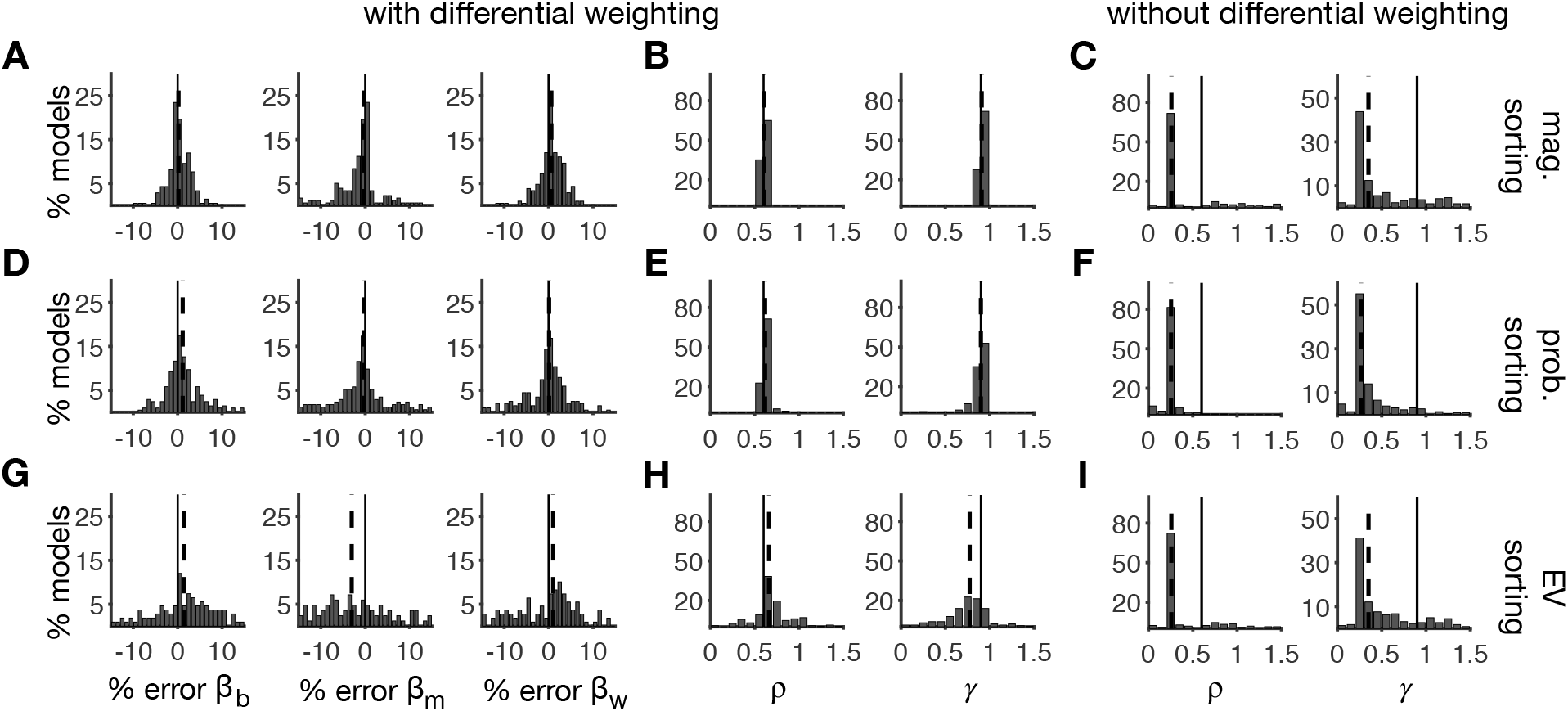
The model with differential weighting can estimate original parameters without a significant bias, whereas the model without differential weighting systematically underestimates *ρ* and *γ* for the data generated with differential weighting and nonlinear utility and probability weighting functions. (**A**) Plots show the distribution of the relative error in the estimation of weights assigned to the best, medium, and worst outcomes (from left to right). For these simulations, the values of individual gambles were computed using nonlinear utility and probability weighting functions (*ρ* = 0.6 and *γ* = 0.9), and outcomes were sorted based on reward magnitude. (**B**) The distribution of estimated values for *ρ* and *γ* across all models. Overall, the estimated parameters were centered around the actual values. (**C**) The distribution of estimated values for *ρ* and *γ* when choice data were fit with a model without differential weighting. The estimated parameters were significantly smaller than the actual values. (**D-F**) The same in panels A-C but for sorting based on reward probability. (**G-I**) The same as in panels A-C but for sorting based on *EV*. There was a small systematic bias in estimated parameters when the choice data were generated with sorting based on *EV.* Nevertheless, this bias does not affect our results since most of our subjects sorted outcomes based on reward magnitude or probability (and not *EV*).

Finally, to compare our model and competing models for valuation of complex gambles, we used four rank-dependent models to fit our experimental data. This includes cumulative prospect theory (CPT), transfer of attention exchange (TAX), decision field theory (DFT) and salience theory of choice (STC) (see Materials and Methods). We found that our model can better predict choice behavior for the majority (55%) of subjects (**Fig. 5A**). We also compared the ability of our model versus each of the competing models and found that the best competing model could fit data better for only 1/3 of subjects (percentage of subjects that were better fit with a competing model: CPT = 14%; TAX = 20%; DFT = 12%; STC = 32.8%; **Fig. 5B**).

**Figure 5.**
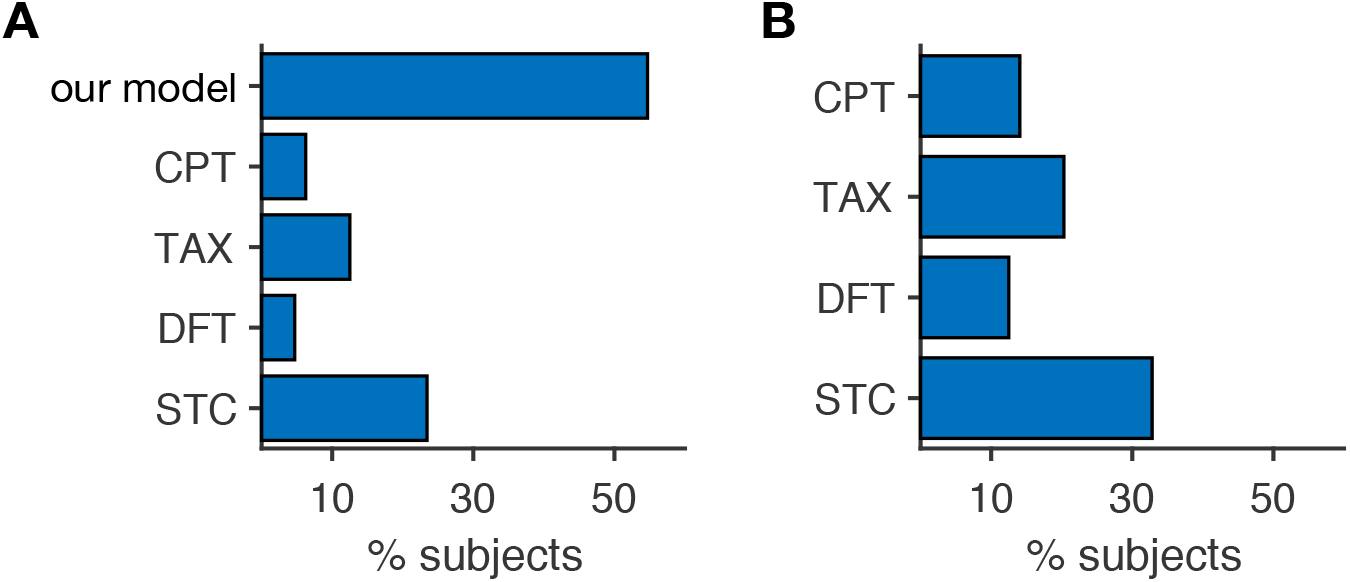
Our model accounts for choice behavior in the majority of subjects. (**A**) Plotted is the percentage of subjects whose choice behavior was best fit by our model and the four competing models. (**B**) Plotted is the percentage of subjects whose choice behavior was better fit by one of the four competing models (versus our model).

### Subjects assigned larger weights to the best and worst outcomes in terms of reward magnitude and probability

Having established that most subjects used differential weighting to combine reward values of possible outcomes, we then examined the weights assigned to the three outcomes within a gamble. For subjects who sorted outcomes based on a single reward attribute, we observed significant differences in weight assignments (**Fig. 6A–D**). More specifically, the weight assigned to the best (largest magnitude or probability) outcome was significantly greater than that of the middle outcome (*β_b_* − *β_m_* = 0.13±0.27 (mean±std), two-sided sign-test, *P* = 0.02, *d* = 0.44; **Fig. 6C**). The worst outcome also had a significantly greater weight compared to the middle outcome (*β_w_* − *β_m_* = 0.09±0.37 (mean±std), two-sided sign-test, *P* = 0.0005, *d* = 0.23; **Fig. 6D**). There was no significant difference between the weights for the best and worst outcomes (two-sided sign-test, *P* = 0.61, *d* = 0.11). These results illustrate that the most important outcomes (best and worst based on a given subject’s sorting) were assigned larger weights for the construction of overall reward value. In contrast, for subjects who sorted outcomes based on an integrated value (*EV, EV* with *w*(*p*), *EU*, or *SU*), there were no significant weight differences between the best and middle outcomes (*β_b_* − *β_m_* = 0.06±0.30 (mean±std), twosided sign-test, *P* = 0.7, *d* = 0.17; **Fig. 6E–H**) or between the worst and middle outcomes (*β_w_* − *β_m_* = −0.08±0.45 (mean±std), two-sided sign-test, *P* = 0.98, *d* = 0.19). These results indicate that differential weighting of gamble outcomes was consistent mainly among subjects who sorted outcomes based on a single piece of reward information.

**Figure 6.**
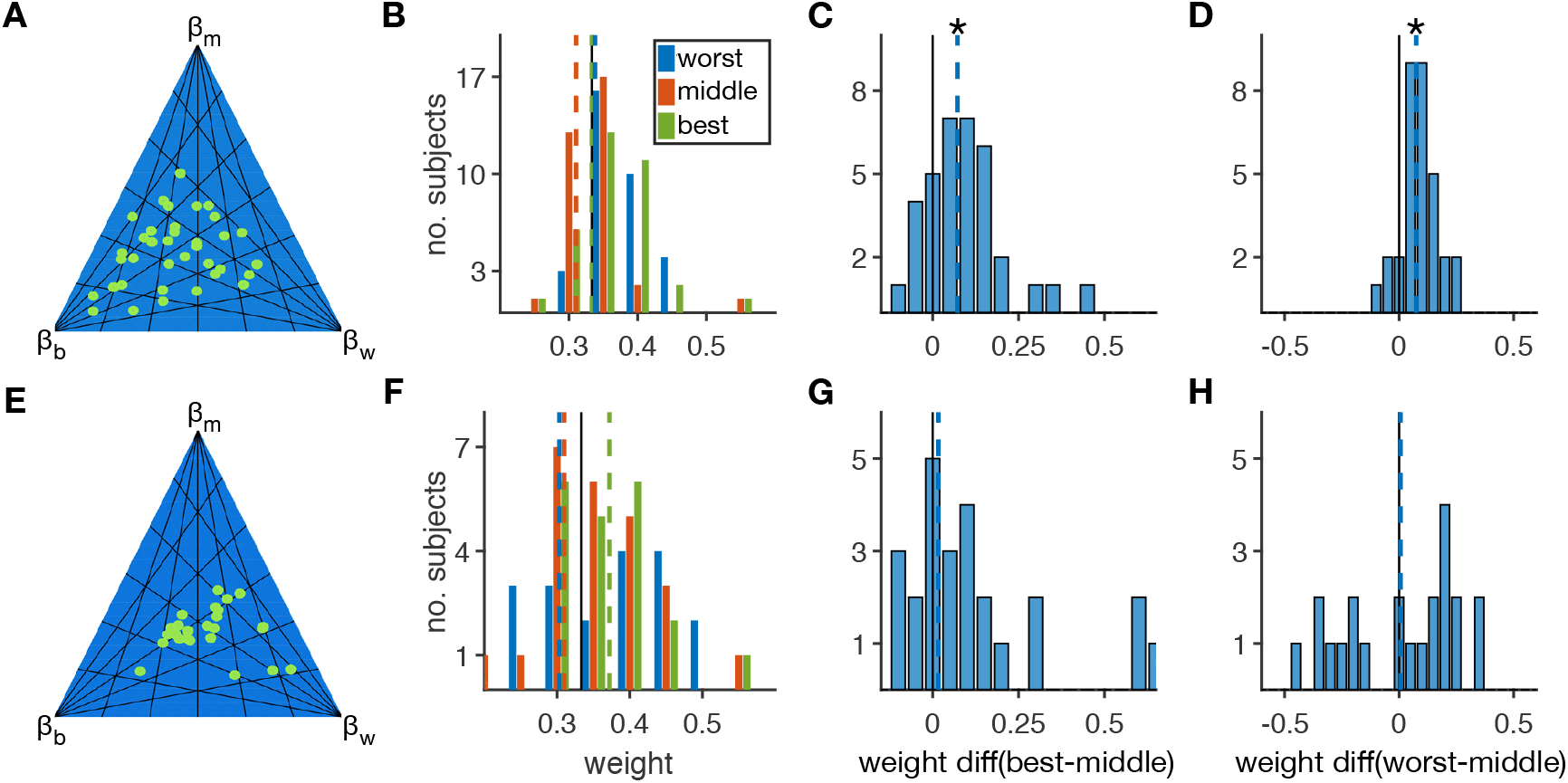
Differential weighting was consistent among subjects who sorted outcomes based on either reward magnitude or probability. (**A**) The ternary plot shows the values of differential weighting for subjects who used a single gamble attribute to sort possible outcomes (*N* = 39). Each dot indicates a set of weights indicated by the distance of the dot from the three corners. (**B**) Plot shows the distribution of absolute weight values assigned to each outcome within a three-outcome gamble. Dashed lines represent median weights. The median weights assigned to the best, middle, and worst outcomes (as determined by either reward magnitude or probability) were 0.33, 0.31, and 0.34, respectively. (**C**) Plot shows the distribution of relative weight differences between the best and middle outcomes. The relative weights were computed by normalizing the two weights by their sum. Dashed lines represent the median, and the asterisk indicates that the median difference was statistically different than zero (two-sided sign-test, *P* < 0.05) (**D**) Plot shows the distribution of relative weight differences between the worst and middle outcomes. Conventions are the same as in panel (**C**). (**E–H**) Plots are similar to panels **A–D** but show the results for subjects who used an integrated value (*EV*, *EV* with *w*(*p*), *EU*, or *SU*) for sorting and assigning weights (*N* = 25).

### Differential weighting enabled subjects to more easily and quickly choose between complex gambles

To address possible advantages of differential weighting, we examined the influence of this mechanism on overall risk preference and whether it allowed subjects to make decisions more easily. Critically, we designed the complex-gamble task such that the pair of gambles presented on each trial has similar *SU* (using a wide range of reward probabilities) in order to detect additional mechanisms involved in the construction of reward value (see Materials and Methods). To quantify the influence of differential weighting on overall risk preference, we computed the relative change in the value of each gamble after the inclusion of differential weighting, for a given set of weights (for all gambles used in the complex-gamble task). To quantify how easily a given subject could distinguish between pairs of gambles, we defined “discriminability” based on the subjective values of gambles estimated for that subject (see Materials and Methods). We computed discriminability for each subject in the simple-gamble task as well as in the complex-gamble task using the best subject-specific models with and without differential weighting.

The influence of differential weighting on risk preference depended on which outcome was assigned the largest weight. The overall value of a three-outcome gamble always increased if the outcome with the highest expected value (or subjective utility) was assigned with the largest weight, resulting in more risk-seeking behavior (**Fig. 7-1**). In contrast, the overall value decreased if the outcome with the lowest expected value was assigned the largest weight, resulting in more risk-aversive behavior. The change in subjective value due to differential weighting was more complex in the case of sorting based on reward magnitude, but on average, assigning a larger weight to the best outcome (i.e. outcome with the largest magnitude) resulted in an increase in the value of gambles and thus increased risk-seeking behavior (**Fig. 7A**). This was the case for many subjects who sorted gamble outcomes based on reward magnitude. Similar but weaker changes in subjective values were observed in the case of sorting based on reward probability (**Fig. 7B**).

**Figure 7.**
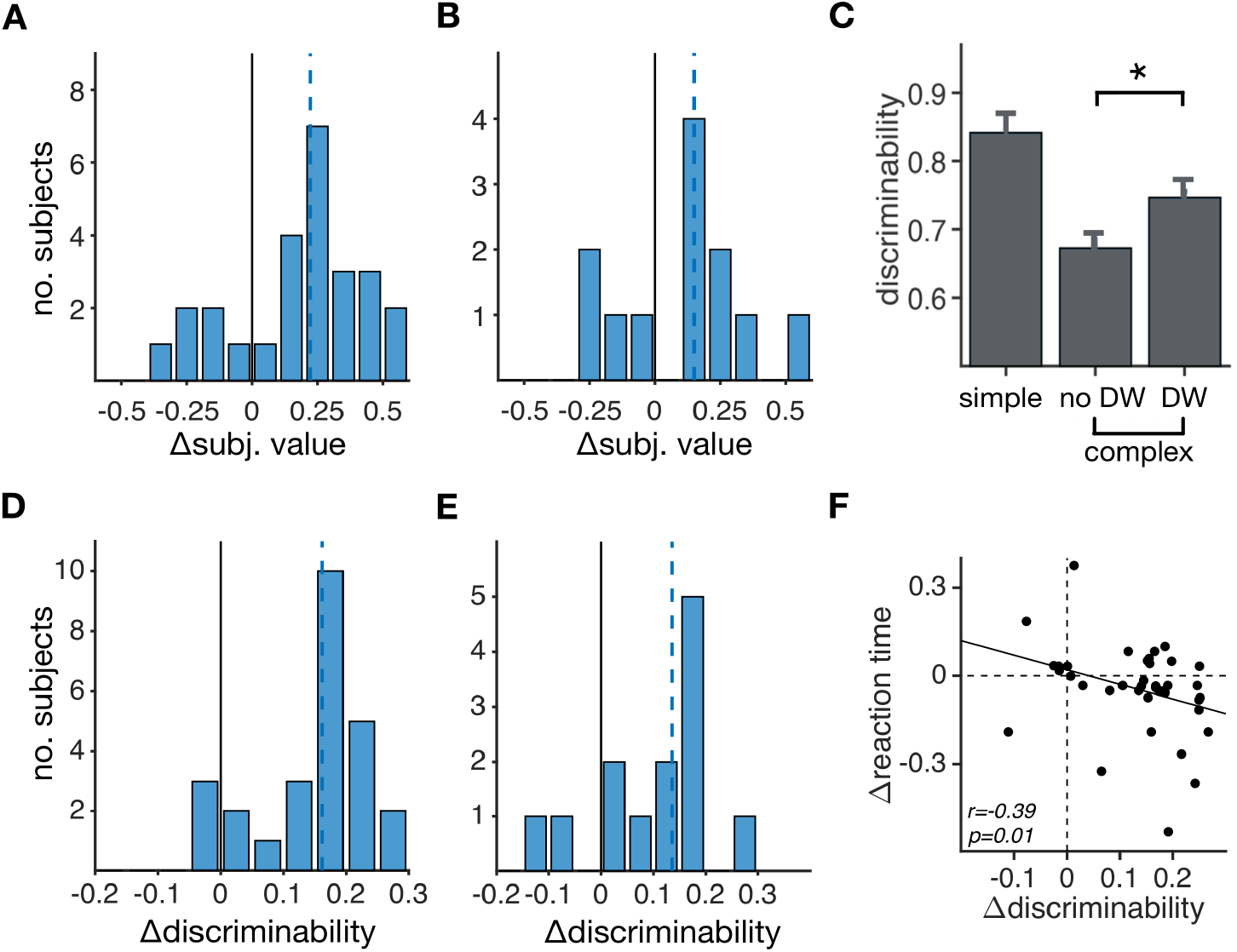
Differential weighting enabled subjects to make decisions more easily and quickly. (**A-B**) Influence of differential weighting on risk preferences. Plots show the distributions of changes in subjective value of a three-outcome gamble after including differential weighting with a given set of weights, using either reward magnitude (A) or probability (B) for sorting. Dashed lines represent the median, which were equal to 0.19 and 0.16 in A and B, respectively. (**C**) Plot shows the discriminability of subjects for the simple-gamble (simple) task as well as the complex-gamble task based on the models with and without differential weighting (DW). The inclusion of differential weighting significantly increased discriminability in the complex-gamble task (two-sided sign-test, *P* = 1.4×10”^10^, *d* = 0.51). (**D-E**) Influence of differential weighting on discriminability. Plots show discriminability for a given set of weights, as indicated by their distances from each corner of the ternary plot, and using magnitude (D) or probability (E) for sorting. Other conventions are similar to those in panels A-B. (**F**) Plotted is the change in median reaction time between the simple- and complex-gamble tasks versus the corresponding change in discriminability due to differential weighting for individual subjects.

**Figure 7-1.**
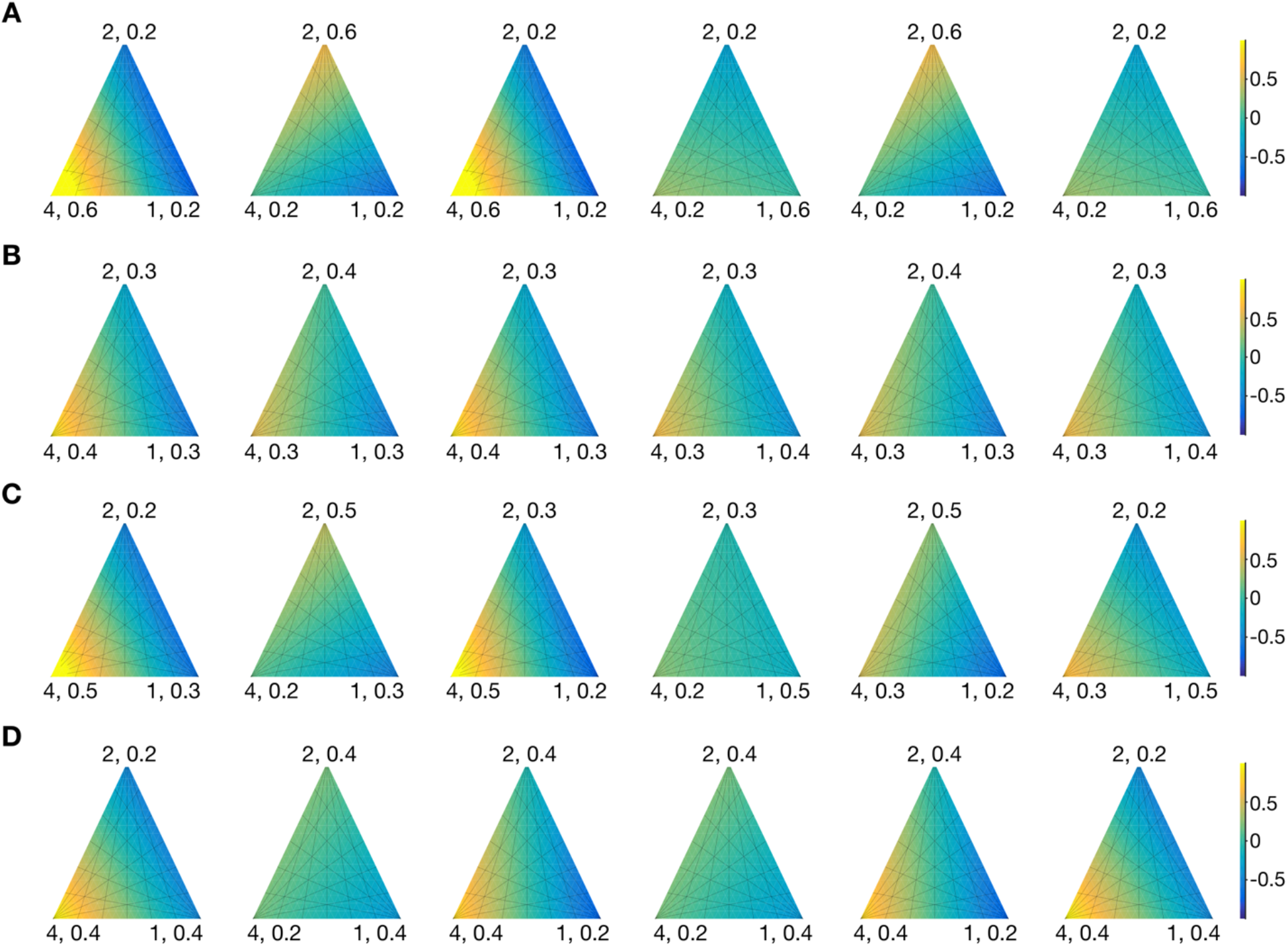
Changes in risk preference due to differential weighting. (**A**) Plots show the change in the subjective value of a three-outcome gamble after including differential weighting with a given set of weights indicated by their distances from each corner of the ternary plot and using magnitude for sorting. The two values on each corner are the reward magnitude and probability associated with the best (lower right corner), medium (upper corner), and worst (lower left corner) outcomes, as determined by reward magnitude. The subjective value of the gamble increased when the largest weight was assigned to the outcome with the largest expected value (i.e. the product of the reward magnitude and probability on a given corner). (**B-D**) The same as in panel A but with different sets of reward probabilities associated with different reward magnitudes.

The effect of differential weighting on discriminability was more complex, but overall, differential weighting increased discriminability for most weight values (**Fig. 7D-E**). As expected, discriminability was smaller across all subjects in the complex-gamble task compared to the simple-gamble task because of differences in task design (**Fig. 7C**). The inclusion of differential weighting, however, significantly increased discriminability across all subjects during the complex-gamble task (two-sided sign-test, *P* = 1.4×10^−10^, *d* = 0.51). In other words, subjects who used differential weighting could more easily discriminate and thus, choose between gambles.

To test whether this increase in discriminability also allowed for faster decision making, we examined the correlation between the change in the average reaction time between the simple- and complex-gamble tasks and the corresponding change in discriminability due to differential weighting within individual subjects. We found a significant negative correlation between the change in reaction time and the change in discriminability for subjects who sorted outcomes based on reward probability or magnitude (Pearson correlation: *r* = −0.32, *P* = 0.035; Spearman correlation, *r* = −0.39, *P* = 0.017; **Fig. 7F**). A similar result was found when considering all subjects (Pearson correlation: *r* = −0.34, *P* = 0.006; Spearman correlation, *r* = −0.38, *P* = 0.002). This indicates that subjects became relatively faster in the complex-gamble task depending on the extent to which they used differential weighting for discrimination between gambles. Together, these results demonstrate that differential weighting enabled subjects to more easily and quickly select between three-outcome gambles.

## Discussion

Here, we examined how valuation and decision-making processes depend on the complexity of risky options by comparing choice between simple and three-outcome gambles within the same individuals. We found that choice between simple gambles was consistent with PT since evaluation based on subjective utility provided the best fit to our data. When evaluating three-outcome gambles, subjects also combined reward probability and magnitude to assign a value to each gamble outcome, but at the same time, most subjects differentially weighted possible outcomes based on either reward magnitude or probability. This heuristic weighting of possible outcomes, in turn, allows for a more dynamic construction of reward value and enables easier and faster decision making, especially for difficult choices between options with similar objective or subjective values. Together, our results reveal a plausible, salience-driven mechanism (perhaps via attentional deployment) underlying value construction for complex gambles.

Currently, there are a number of sophisticated models for valuation and choice between complex gambles. Similar to our model, most of these models rely on a rank-dependent mechanism for processing alternative outcomes, and our model can be considered as a special case of the STC model. Unlike our model, however, these models rely on complex computations. In addition, none of these complex models have been used to fit choice data from individual subjects, and thus, there is no evidence that they can capture individual variability. We find that although choices of a small fraction of subjects are better fit by more complex models, our simple model provides a better fit for most individual subjects.

Given our limited processing resources, exhaustively weighting and summating all possible outcomes to evaluate an option is not feasible unless we can simplify valuation and decision-making processes using some form of heuristics (Brandstätter et al., 2006; Gigerenzer and Goldstein, 1996; Gigerenzer and Gaissmaier, 2011). In many cases, information must somehow be prioritized in order to avoid cognitive overload. Such prioritization has been assumed to be performed mainly via attentional mechanisms (Klein, 1988; Treisman and Gelade, 1980; McLeod et al., 1988; Russell and Kunar, 2012; Wolfe and Horowitz, 2004; Watson and Kunar, 2010). It has been shown that both bottom-up and top-down attention can influence processing and integration of reward information and ultimately choice behavior (Krajbich et al., 2010; Tsetsos et al., 2012; Kunar et al., 2017). By selectively processing certain outcomes within complex risky options, the decision maker can reduce computational demands required for evaluation of such options.

Heuristics similar to those included in our model have been proposed also for reducing the computational demands and complexity of valuation and decision processes, such as when choosing between multiple alternatives or during multi-attribute choice even in the absence of risk (Payne et al., 1993; Gigerenzer and Goldstein, 1996; Gigerenzer and Todd, 1999). In the case of multi-attribute options, decision making can be very difficult because it requires weighting the pros and cons of options that differ across multiple, sometimes incommensurate, dimensions (Fellows, 2006). It has been shown that in such situations, humans use various strategies and heuristics such as differential weighting of different dimensions, limiting the amount of information to be processed, and reducing the number of alternatives to be considered (Payne et al., 1993; Gigerenzer and Todd, 1999). Although we assumed that each gamble is assigned with an overall subjective value, valuation and choice processes in our model also can be interpreted as weighted comparisons across individual outcomes based on reward probability or magnitude; that is, the subjects could directly choose between gambles by comparing values for similar individual outcomes and combine such comparisons across all possible outcomes (Tversky, 1969, 1972).

Critically, we observed a dissociation between what drives the evaluation of outcome value (integrated value) and what drives the weighting process (single gamble attribute), which indicates that selective processing of reward information may not rely on both probability and magnitude. The sorting of outcomes based on a single attribute can be seen as a more general case of the elimination by aspect theory (Tversky, 1972). One possible mechanism for such processing is selective attention (Hayden et al., 2008, 2007; Ludvig et al., 2014; Shimojo et al., 2003; Busemeyer and Townsend, 1993; Roe et al., 2001). In our task, the most relevant form of attention is feature-based attention, which could selectively enhance the representation of certain visual attributes (e.g., color or size) at the expense of the others (Carrasco, 2011). Given the visual presentation of gambles in our experiment, subjects could attend to certain “learned” reward features (color and size) within the gamble and thus, weigh outcomes by their reward salience. Interestingly, in our experiment, subjects assigned larger weights to both the best and worst outcomes, which indicates that extreme outcomes are most salient. A plausible neural mechanism for this differential weighting could be biased competition associated with feature-based attention (Reynolds et al., 1999; Kastner and Ungerleider, 2001; Beck and Kastner, 2005) or competition at multiple levels of value representation (Jocham et al., 2012; Hunt et al., 2014). The increased weighting of the best outcome in terms of magnitude is consistent with observed increases in attention toward larger rewards (Raymond and O’Brien, 2009; Della Libera and Chelazzi, 2006) and results in risk-seeking behavior, whereas larger weighting of the worst outcome can contribute to risk-aversion. Taken together, our results suggest that attentional processes could contribute to differential weighting of reward outcomes by their salience in order to simplify the valuation process.

The observed dissociation between the type of reward information that contributes to the evaluation of individual outcomes and to the weighting of alternative outcomes highlights the importance of attention for adaptive choice under risk at the expense of optimality (Farashahi et al., 2017a,b). More specifically, although each possible outcome should be evaluated based on a quantity that combines different pieces of reward information (and thus could be optimal), differential weighting based on a single quantity (e.g., reward magnitude or probability) can enable flexibility depending on the state of the decision maker. This dissociation also suggests separate mechanisms through which reward influences choice and selective processing of information (Soltani et al., 2016; Rakhshan et al., 2018). For example, when hungry, the decision maker could attend more to cues that represent the amount of food, or reward. This could result in more risk-seeking behavior but also faster and easier decision making, both of which are crucial for survival. Therefore, differential weighting of outcomes provides a plausible mechanism for flexible risk attitudes (Bruhin et al., 2010; Lattimore et al., 1992; Heyand Orme, 1994; Huber et al., 1982; Ludvig et al., 2013; Rigoli et al., 2015; Stewart et al., 2003; Fujimoto and Takahashi, 2016) and results in a tradeoff between optimality and flexibility. We have recently shown that a simpler version of our model can account for monkey’s choice behavior during choice between simple gambles, providing further evidence for differential weighting (Farashahi et al., 2018). We speculate that observed deviations from normative theories of choice could be due to such weighting mechanisms, reflecting the flexibility required for decision making in dynamic environments (Payne et al., 1988). Together, our results shed light on neural mechanisms of choice under risk in naturalistic settings, and moreover, highlight the role of attention in the flexible construction of reward value for complex gambles.

## Author Contributions

A.S, E.C, and M.S. conceptualized the project, designed the experiments, analyzed and interpreted the data. E.C. performed experiments and acquired the data. A.S. and M.S designed the models. M.S. implemented the models and preformed the simulations. A.S. and M.S. analyzed model simulations. A.S. and E.C. wrote the main manuscript text, which all authors have reviewed. All authors approved the final version of the manuscript. Study supervision was performed by A.S.

## Acknowledgements

We would like to thank Daeyeol Lee for helpful comments on the manuscript. This work is supported by NSF grant (EPSCoR Award # 1632738) to AS.

